# Ontology-Guided Pathway Activity Identifies a Cell-Intrinsic Defense Response Program Associated with MEK Inhibitor Sensitivity

**DOI:** 10.64898/2026.07.25.740732

**Authors:** Chimdi Walter Ndubuisi, Project Cancer Drug Response Collaboration

**Affiliations:** University of Missouri – Columbia

## Abstract

Predicting cancer drug response from gene expression requires models that expose which biological pathways drive cell-line-specific sensitivity. We introduce Gene-Ontology Pathway Attention (GOPA), whose core module computes deterministic attention weights softmax(*x_c_* · *B̂*) from expression and the column-normalized gene-term annotation matrix with no learned parameters. Applying GOPA to 542 drugs across the GDSC panel under leave-cell-line-out evaluation, we find that defense response pathways predict sensitivity to kinase inhibitors in GDSC, with the strongest and most confound-resistant signal in MEK/MAPK inhibitors. This association shows cross-assay support in PRISM (11 of 11 overlapping drugs; all *p <* 0.002) and retains 72% signal strength after controlling for five confounds, but was not reproduced in the gCSI panel (different response metric, smaller sample, MEK inhibitors absent), indicating the finding may be MAPK-pathway-specific and assay-dependent. On the 16-drug benchmark, XGBoost achieves the lowest RMSE (1.185); GOPA (1.226) is the strongest neural model. On the full 542-drug panel, target-encoded XGBoost matches GOPA on RMSE (1.322 vs. 1.327); GOPA achieves higher residual Pearson correlation (0.469; 95% CI [0.453, 0.485]) than target-encoded XGBoost (0.391; [0.379, 0.404]); paired difference +0.078 [0.062, 0.093]; GOPA wins on 363 of 539 drugs (67.3%); Wilcoxon *p* = 1.1 × 10^−23^). When pathway representations are evaluated with matched downstream learners, simple gene-set projections achieve equivalent prediction, indicating that GOPA’s value lies in its deterministic, population-comparable pathway summaries rather than representational superiority. A controlled geometry comparison shows Poincaré-ball embeddings preserve GO graph distances better than Euclidean (*ρ* = 0.732 vs. 0.474) while Euclidean embeddings achieve stronger ancestor retrieval; neither geometry improves prediction (ΔRMSE = +0.008; *p* = 0.18).

## 1 Introduction

Geometric deep learning has achieved notable success on molecular graphs, protein structures, and knowledge graphs [Bronstein et al., 2021], yet biological ontologies remain underexploited as geometric objects. The Gene Ontology (GO) organizes gene function into a DAG spanning *>*40,000 terms [Gene Ontology Consortium, 2023]. Its tree-like branching structure makes it a natural candidate for hyperbolic representation learning, where exponential volume growth matches the exponential branching of hierarchies [Nickel and Kiela, 2017, Ganea et al., 2018, Chami et al., 2019]. Two questions remain open: does geometric fidelity to the ontology improve downstream task performance, and can ontology-aware architectures surface biology that post-hoc interpretation does not?

Cancer drug-response prediction provides a rigorous testbed. Data leakage from repeated measurements inflates apparent gains when models interpolate identity effects [Barretina et al., 2012, Iorio et al., 2016, Hostallero et al., 2024], and even accurate models rarely expose *which pathways* drive sensitivity. Prior work has used GO structure as an architectural prior: DCell and Drug-Cell organize network layers to mirror the GO hierarchy [Ma et al., 2018, Kuenzi et al., 2020], but these approaches hardwire topology without sample-specific pathway representations. Graph-based models including TGSA [Zhu et al., 2022], GraphDRP [Nguyen et al., 2022], and DeepTTC [Jiang et al., 2022] use protein-protein interaction or molecular graphs rather than the functional ontology.

We introduce **Gene-Ontology Pathway Attention (GOPA)**, a geometric deep learning architecture that separates *what pathways are active in a sample* (computed deterministically) from *what those activations mean for prediction* (learned). The attention weights softmax(*x_c_* · *B̂*), where *B̂* is the column-normalized gene-term annotation matrix with ancestor-propagated annotations, contain no learned parameters. This provides structurally grounded interpretive properties: the weights are invariant to training, biologically grounded through GO annotations, Lipschitz-stable, and population-comparable (Supplement S27). Pathway attention can therefore be computed for any cell line without model inference, enabling validation across independent datasets and tissue contexts without retraining. Learned parameters enter through GO term embeddings (refined by a GNN on the GO DAG, optionally in Poincaŕe-ball geometry) and gated aggregation that transforms attention-weighted embeddings into predictions. The key contributions are:

1. A **deterministic, parameter-free attention mechanism** (the Gene-Ontology Bridge) that computes sample-specific pathway activity from expression and GO annotations, enabling population-scale biological validation across datasets, functional assays, and tissue contexts without model retraining.
2. **A controlled geometry comparison** of ontology embedding: Poincaŕe-ball embeddings preserve GO graph distances better than Euclidean (*ρ* = 0.732 vs. 0.474) while Euclidean embeddings achieve stronger ancestor retrieval (MRR 0.142 vs. 0.131); neither geometry improves drug response prediction (ΔRMSE = +0.008; *p* = 0.18), localizing the predictive signal to the annotation-based attention mechanism.
3. **Biological discovery**: defense response pathways predict sensitivity to kinase inhibitors in GDSC, strongest for MEK/MAPK inhibitors (partial *ρ <* −0.15 after all-confound correction), with cross-assay PRISM support (11 of 11 overlapping drugs), CRISPR functional association, PD-L1 expression association, and confound-controlled partial correlations (72% retention). The association was not reproduced in gCSI, indicating it may be MAPK-pathway-specific and assay-dependent.
4. **A systematic evaluation** following DrEval-style benchmarking [Hostallero et al., 2024] showing that tree-baseline superiority on RMSE reflects drug-encoding efficiency; with target encoding, trees match GOPA on RMSE while GOPA achieves higher residual Pearson (0.47 vs. 0.39; paired difference +0.08 [0.06, 0.09]; Wilcoxon *p <* 10^−23^). With matched downstream learners, simple pathway projections achieve equivalent prediction, establishing that GOPA’s value lies in interpretability rather than representational superiority.

In this study, we test whether a deterministic projection of expression through GO annotations yields useful and reproducible pathway summaries for cancer drug response. We evaluate the representation on leakage-aware GDSC splits, compare it with matched pathway scores and predictive baselines, test the defense response association across pharmacological and functional datasets, and isolate the effect of Euclidean versus hyperbolic ontology embeddings. The analyses distinguish structural interpretability, predictive dependence, and ontology representation quality rather than treating them as equivalent.

## 2 Related Work

**Ontology-structured neural networks.** DCell [Ma et al., 2018] pioneered using the GO hierarchy as a network architecture, mapping each GO term to a subnetwork whose connectivity mirrors the ontology. DrugCell [Kuenzi et al., 2020] extended this to drug response, hardwiring network topology to the GO DAG. P-NET [Elmarakeby et al., 2021] applied a similar biologically informed architecture to prostate cancer classification. All three approaches embed the ontology into the architecture itself, coupling interpretation to learned parameters. GOPA instead uses the ontology as an attention scaffold: the Gene-Ontology Bridge computes pathway activations deterministically from expression and the annotation matrix, separating interpretation from learned prediction.

**Graph neural networks for drug response.** TGSA [Zhu et al., 2022] uses protein-protein interaction graphs with similarity augmentation; GraphDRP [Nguyen et al., 2022] combines molecular and cell-line graphs; DeepTTC [Jiang et al., 2022] applies transformers to SMILES strings. These methods operate on molecular or interaction graphs rather than the functional ontology, and none provide parameter-free pathway interpretation. DeepCDR [Liu et al., 2020] integrates multiomic data via 1D convolutions. All were benchmarked under protocols that DrEval [Hostallero et al., 2024] showed systematically inflate performance through data leakage.

**Attention, feature attribution, and interpretable pharmacogenomics.** Post-hoc interpretation methods including SHAP [Lundberg and Lee, 2017] and integrated gradients [Sundararajan et al., 2017] provide feature attributions for trained models but produce different pathway rankings depending on the model, hyperparameters, and background distribution. Concerns about learned attention faithfulness [Jain and Wallace, 2019, Wiegreffe and Pinter, 2019] motivate architectures where interpretation is structurally independent of the learned prediction objective. Pathway activity scoring methods including GSVA [Hänzelmann et al., 2013], ssGSEA [Barbie et al., 2009], AUCell [Aibar et al., 2017], and PROGENy [Schubert et al., 2018] provide sample-specific pathway summaries but use curated or statistically derived gene sets rather than ontology-guided projections, and do not integrate with end-to-end prediction architectures.

**Leakage-aware evaluation and pharmacogenomic benchmarking.** DrEval [Hostallero et al., 2024] demonstrated that many published drug response models derive apparent gains from data leakage via shared cell lines or drugs across train/test splits. Our evaluation follows DrEval principles throughout: leave-cell-line-out splitting, drug identity features in all baselines, and explicit reporting of per-drug-mean null performance.

**Gap.** Existing ontology-structured and pathway-informed models either embed ontology structure inside learned parameters, use post-hoc gene attribution, or evaluate prediction without separating ontology representation quality from task utility. Few studies test whether deterministic sample-level pathway projections add value beyond matched pathway scores under leakage-aware evaluation with cross-screen biological validation. GOPA addresses this gap by providing a deterministic, population-comparable pathway representation within an end-to-end prediction architecture, evaluated with matched baselines and cross-screen validation.

## 3 Results

### 3.1 Defense Response Pathway Association with Kinase Inhibitor Sensitivity

Among 420 PRISM-validated novel pathway associations (1.3–2.9× enrichment over correlation-preserving permutation nulls; *p <* 0.001), a convergent pattern emerges (Figure 1): defense response pathways (GO:0006952, GO:0006955, GO:0042110) predict sensitivity across 12 kinase inhibitors from 5 target families: EGFR (afatinib *ρ*_PRISM_ = −0.303, gefitinib −0.238, osimertinib −0.247, pelitinib −0.291), MEK (trametinib −0.298, selumetinib −0.156, refametinib −0.193, PD0325901), SRC/ABL (bosutinib −0.270, dasatinib −0.257), BTK (ibrutinib −0.256), and HER2 (lapatinib −0.270). All show negative *ρ* (higher defense response expression → greater sensitivity). These associations are not annotated in any drug’s target pathway. In GDSC and PRISM, all 12 drugs show the same direction (all *p <* 0.002 for 11 of 11 PRISM-overlapping drugs). The effect is strongest for MEK inhibitors (partial *ρ <* −0.15 after all-confound correction), consistent with the MAPK/PD-L1 mechanistic model. EGFR inhibitors show weaker effects. The defense-response pattern was identified through the intersection of GDSC pathway analysis and PRISM transfer validation; confound and tissue analyses were conducted after pattern identification. PRISM therefore provides cross-assay pharmacological support rather than a formally locked replication. A locked test on gCSI (Section 3.6) did not reproduce the consistent association, indicating the finding may be MAPK-pathway-specific or assay-dependent.

**PD-L1 (CD274) association.** CD274 expression analysis shows an association consistent with MAPK-driven PD-L1 upregulation [Stutvoet et al., 2019]: kinase inhibitors show more negative CD274-response correlations than 530 other drugs (Mann-Whitney *p* = 0.003; rank-biserial *r* = 0.50). The effect is driven by MEK inhibitors (Trametinib *ρ* = −0.145, Refametinib −0.144, PD0325901 −0.122, Selumetinib −0.098; all *p <* 10^−4^) and SRC/ABL inhibitors (Dasatinib *ρ* = −0.191, *p <* 0.002). EGFR inhibitors show weaker or mixed effects. Control drug classes show the opposite direction: HDAC inhibitors have strong positive CD274 correlation (Vorinostat *ρ* = +0.299, Panobinostat +0.290; *p <* 10^−13^), consistent with known HDAC-mediated PD-L1 upregulation.

**Confound control.** Partial Spearman correlations controlling simultaneously for five potential confounds (IFN/JAK-STAT response, cell proliferation, HLA class I expression, epithelial-mesenchymal state, and CD274 expression) show the defense response signal retains 72% of its strength (partial *ρ* = −0.104 vs. raw −0.144; 11/12 drugs negative). IFN/JAK-STAT reduces the signal most (46% retention), consistent with shared interferon signaling. EMT barely affects it (102% retention), confirming orthogonality to epithelial-mesenchymal state. MEK inhibitors retain the strongest confound-resistant signal: Refametinib partial *ρ* = −0.204 (*p <* 10^−13^), Trametinib −0.181 (*p <* 10^−6^) after all-confound correction.

The signal also survives within individual tissues: Lung (*ρ* = −0.41, *p <* 10^−6^, *n* = 134), Breast (−0.47, *p <* 0.001), CNS (−0.43, *p <* 0.01). The Myeloid reversal (*ρ* = +0.46, *p <* 0.01) is consistent with constitutive inflammatory gene activation in hematopoietic lineages. Table 1 consolidates the multi-scale validation evidence for all 12 kinase drugs.

**GO term count robustness.** Across 256, 512, 1024, and 2048 GO terms, 11-12/12 kinase drugs show negative defense-response correlation at every scale (mean *ρ*: −0.089, −0.066, −0.045, −0.067 respectively). The defense response percentile rank remains stable (37th-42nd across scales; Supplement S23).

**Table 1:**
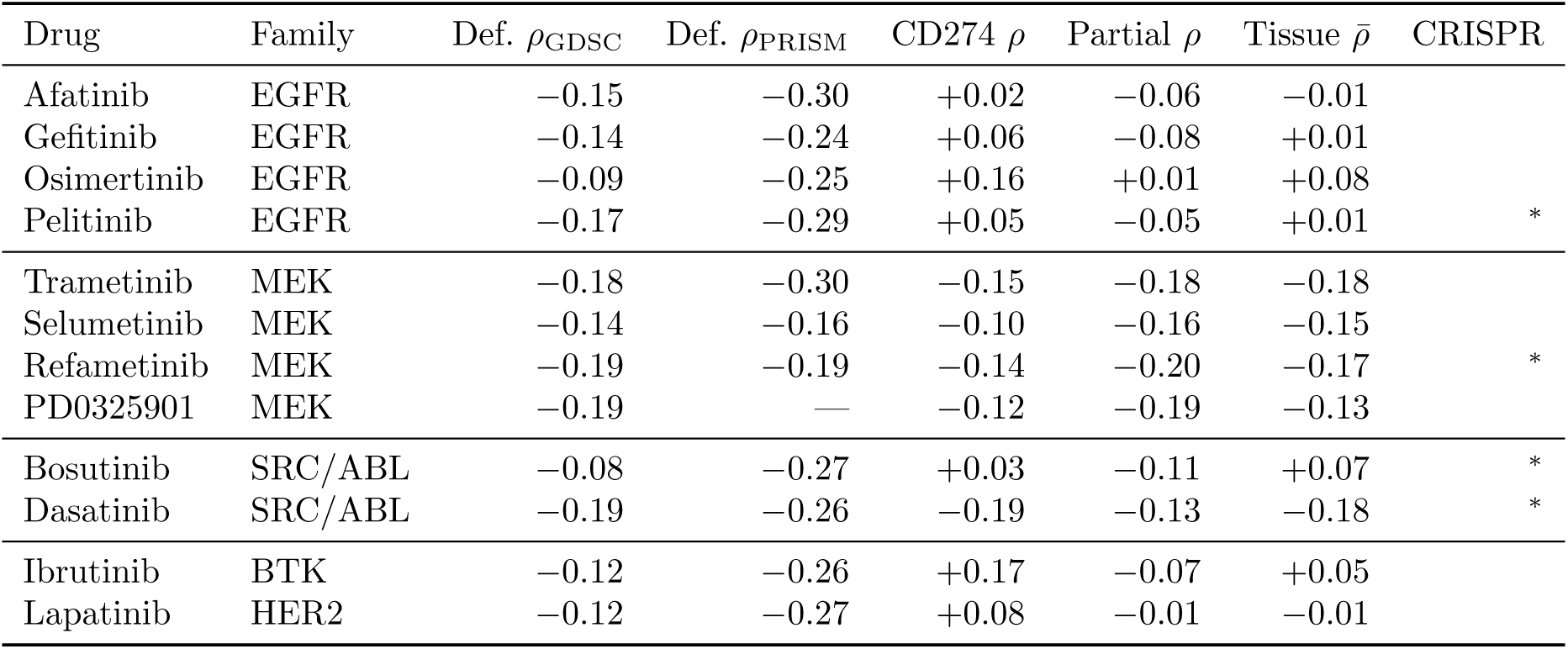
Consolidated validation evidence for the immune–kinase inhibitor sensitivity axis. Defense response (GO:0006952) attention–response Spearman *ρ* in GDSC and PRISM (negative = higher immune expression → greater sensitivity). CD274 *ρ*: PD-L1 expression–response correlation. Partial *ρ*: defense response *ρ* after controlling for IFN/JAK-STAT, proliferation, HLA-I, EMT, and CD274 simultaneously. Tissue *ρ*: mean within-tissue defense response 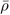 (tissue-of-origin confound removed). CRISPR: top-attention pathway genes show differential essentiality between drug-sensitive and drug-resistant cell lines (^∗^*p <* 0.05).

**Shuffled-annotation control.** Shuffling the annotation matrix *B* column-wise (100 permutations) abolishes the signal: mean *ρ* = +0.019 ± 0.061 vs. −0.187 with real annotations (empirical *p <* 0.01; Supplement S26).

**Representation independence of the discovery.** To test whether the defense-response axis is specific to softmax attention or recoverable by any GO-based representation, we compute defense-response scores using six methods on the full 542-drug panel. All real GO representations recover the axis: column-normalized *x* · *B̂* (mean kinase *ρ* = −0.156; 12/12 drugs negative; 10/12 significant), raw *x* · *B* (−0.135; 12/12 negative), z-scored mean (−0.146; 12/12 negative), and softmax attention (−0.192; 12/12 negative; 12/12 significant). Random gene sets (+0.015; 4/12 negative) and shuffled annotations (−0.008; 5/12 negative) fail to recover the pattern. Softmax attention provides the strongest signal (descriptively 23% larger mean |*ρ*| than column-normalized projection, without paired uncertainty) and the best kinase-specific discrimination: for negative-control HDAC inhibitors, softmax shows near-zero correlation (*ρ* = −0.021, not significant) while simple projections show significant negative correlation (*ρ* = −0.153), indicating that softmax concentration showed stronger descriptive kinase-versus-HDAC separation in this analysis (paired uncertainty for this comparison was not computed). The defense-response discovery is therefore an ontology-projected pathway finding that any real GO representation recovers, with softmax attention providing a modestly stronger and more discriminative signal.

**Multi-axis attention landscape.** PCA on the 542 × 512 drug-GO-term correlation matrix reveals that the attention landscape decomposes into orthogonal biological axes (Supplement S28). The dominant axis (PC1, 51% variance) separates kinase/signaling drugs from chromatin/epigenetic drugs based on cell adhesion and epithelial state: focal adhesion attention predicts resistance for chromatin drugs (*ρ̄* = +0.29) but sensitivity for MEK inhibitors (Refametinib *ρ* = −0.22, Trametinib −0.21; both PRISM-validated). The ERK-chromatin separation is large (Cohen’s *d* = 1.56; *p* = 1.2 × 10^−9^). The defense-response pattern loads primarily on PC3 (8% variance), indicating it is distinct from the dominant adhesion/chromatin axis within this linear correlation-space decomposition (principal components are orthogonal by construction; this does not prove biological independence). The recovery of known biology (EMT axis on PC1) alongside the defense-response pattern (PC3) shows that the pathway projection captures a structured, multi-dimensional landscape.

**Figure 1:**
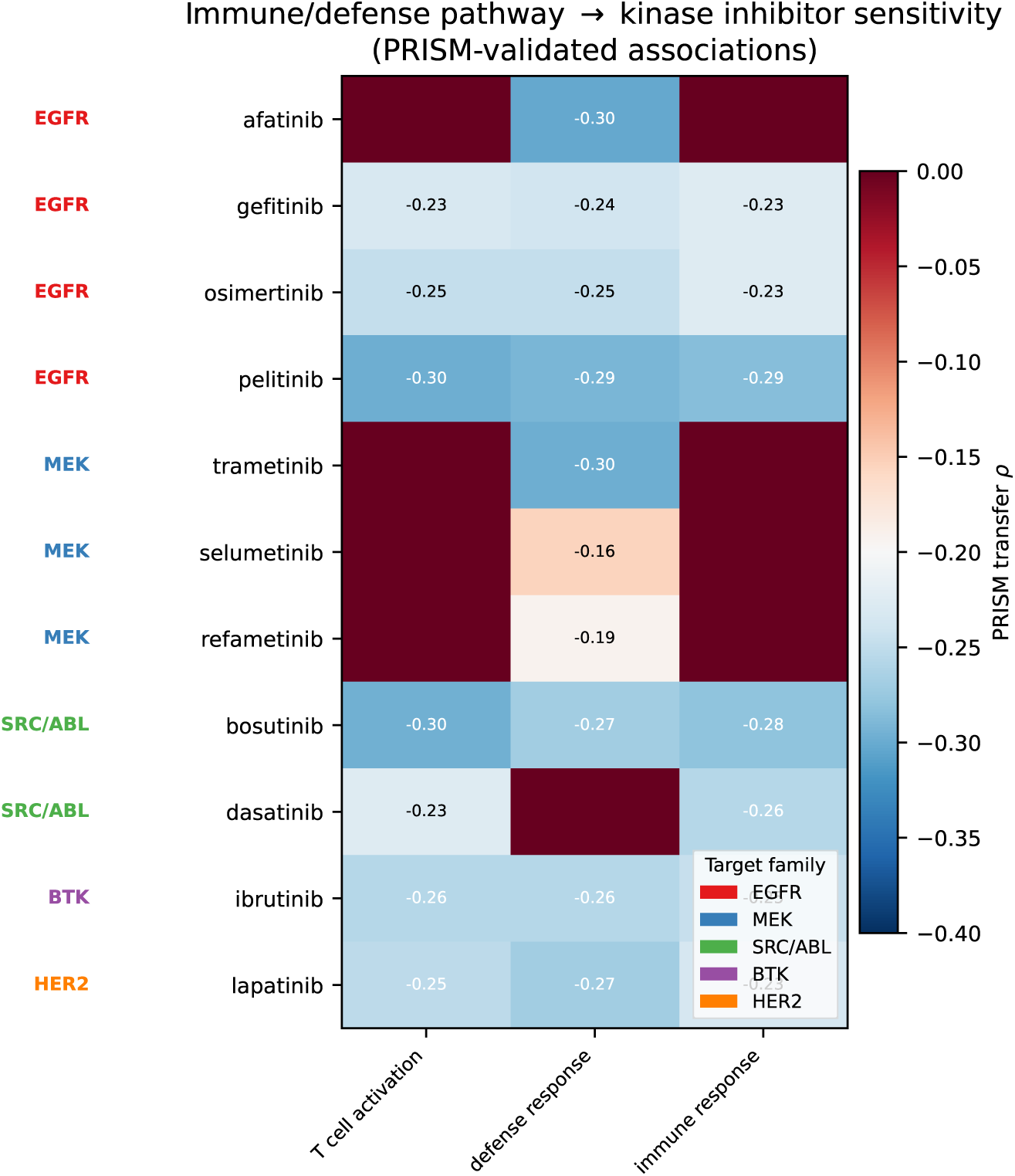
Defense response pathway association with kinase inhibitor sensitivity. Heatmap shows PRISM-validated transfer *ρ* for defense-response GO terms across kinase inhibitors from 5 target families. All associations are negative (higher defense response expression → greater sensitivity) with cross-assay support in PRISM (all *p <* 0.002 for 11 of 11 overlapping drugs); a locked test on gCSI did not replicate the association (see text).

### 3.2 Pathway Validation

GOPA’s attention weights are a deterministic function softmax(*x_c_* · *B̂*) of expression and annotation structure, containing no learned parameters. This makes attention computable for any cell line without model training.

**Figure 2:**
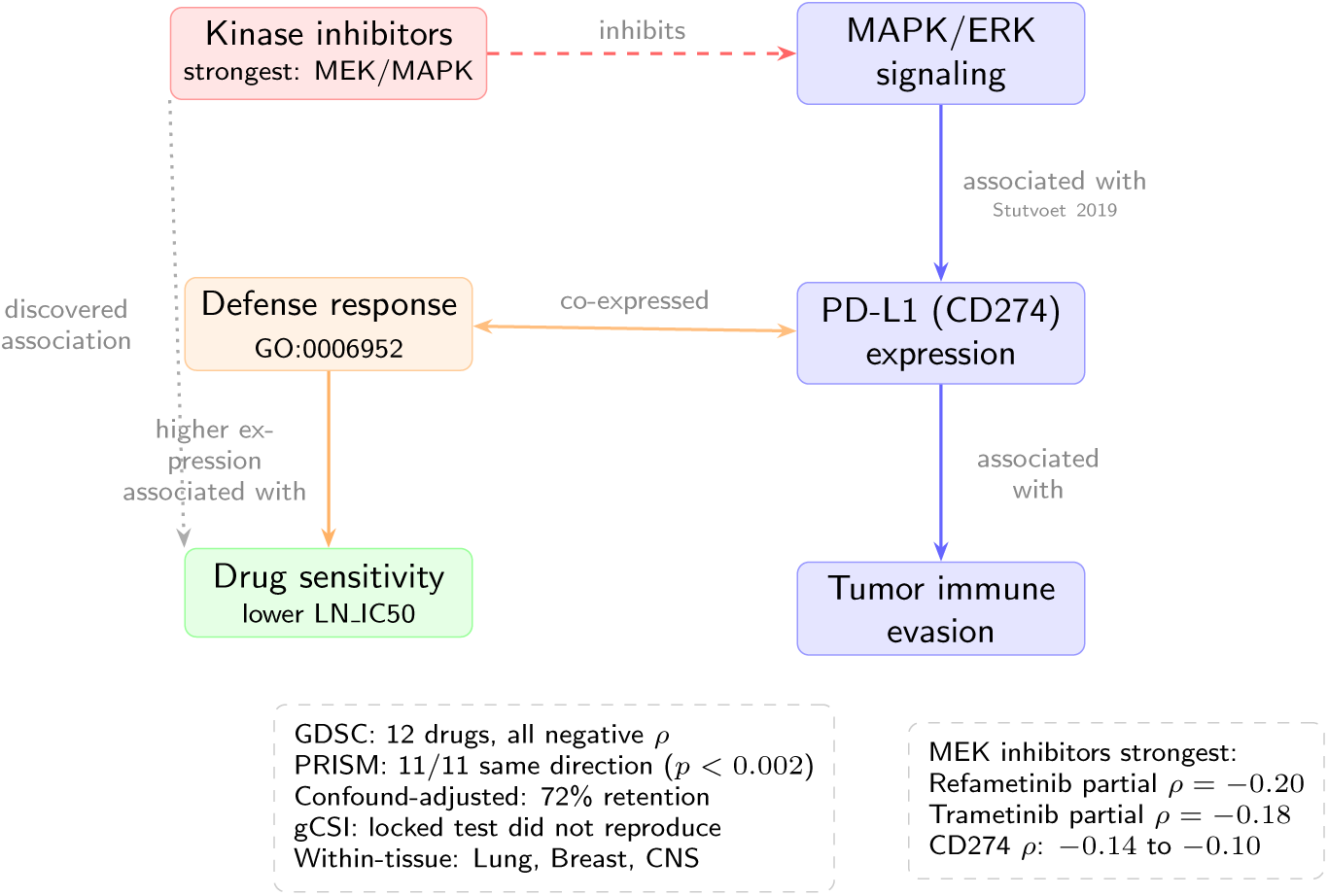
Proposed model linking defense response pathway activation to kinase inhibitor sensitivity. MAPK/ERK signaling is associated with PD-L1 expression in tumor cells [Stutvoet et al., 2019]. Cell lines with high defense response expression are preferentially sensitized by kinase inhibition, with the association strongest for MEK inhibitors. Because these are cell-line models without immune infiltration, the signal reflects tumor-intrinsic defense response programs. Evidence boxes summarize multi-scale validation (Table 1). All arrows represent associations or hypothesized links, not demonstrated causal relationships.

**Case studies (***n* = 6). For drugs with known mechanisms, response-correlated attention aligns with expected biology (Figure 3): MEK inhibitors (PD0325901, Trametinib) show strongest correlation with cytokine-mediated signaling and inflammatory response (*ρ* = −0.30 to −0.35), sharing 6/10 top correlated terms despite chemical distinctness. PARP inhibitors (Olaparib, Talazoparib) correlate with vesicle biology and catalytic activity, with DNA repair terms ranking significantly above chance (65th percentile; *p <* 0.05). Drugs sharing a target pathway have highly correlated attention profiles (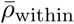 = 0.92 ± 0.02) compared to between-pathway pairs (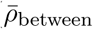 = 0.28 ± 0.27; Mann-Whitney *p* = 0.010).

**Figure 3:**
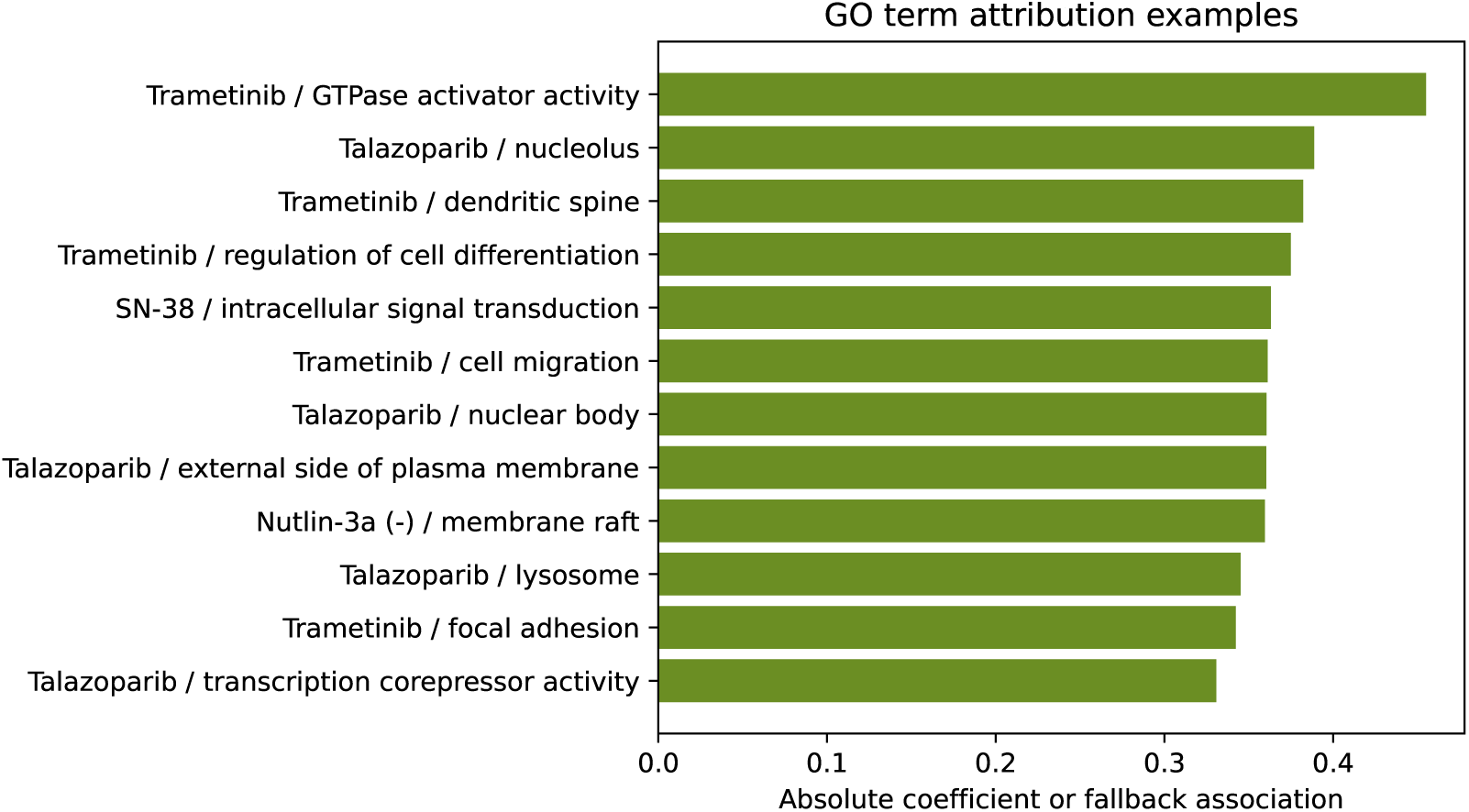
Response-correlated GO term attention for selected drugs.

**Full GDSC panel (***n* = 542). Target-pathway GO terms rank at the 52.4th percentile across 340 drugs with known targets (95% CI [50.8, 54.0]; *p* = 0.006).

**PRISM cross-dataset transfer (158 drugs).** Pathway attention-response correlations transfer from GDSC to the PRISM screen [Corsello et al., 2020] with mean *ρ* = 0.256±0.195 (95% CI [0.225, 0.287]; *t* = 16.5, *p <* 10^−15^; 85.4% individually significant). Of 1,189 novel pathway associations, 420 validate in PRISM (*p <* 0.05, same sign). Under naive independence, 59.5 would be expected; however, GO-term hierarchical overlap and drug correlation inflate this null. Correlation-preserving permutation tests (1,000 permutations) yield more appropriate nulls: drug-label shuffle 172.3 ± 28.5 (2.4× enrichment), GO-term shuffle 321.4 ± 6.3 (1.3× enrichment), and two-way shuffle 144.1 ± 26.3 (2.9× enrichment); all *p* ≤ 0.001 (plus-one permutation estimate, 1,000 permutations). Even under the most conservative null (GO-term shuffle), the observed 420 exceeds the maximum permuted count (353). PRISM shares a substantial cell-line ecosystem with GDSC; this represents cross-assay pharmacological support rather than fully independent cohort validation.

**CRISPR dependency (542 drugs).** Genes in top-attention GO terms show 1.57× greater differential CRISPR essentiality between drug-sensitive and drug-resistant cell lines than random GO term baselines (Wilcoxon *p* = 1.5 × 10^−19^; 203 of 542 drugs significant vs. 124 for bottom-attention controls; Figure 5). Standard DepMap CRISPR dependency measures essentiality without drug treatment; this represents orthogonal functional association, not direct drug-perturbation evidence.

**Leave-tissue-out transfer (5,314 drug-tissue pairs).** Across 11 tissues with ≥100 cell lines, pathway attention signals transfer to held-out tissues with mean *ρ* = 0.145 (drug-level bootstrap 95% CI [0.139, 0.151]; 529 of 542 drugs positive; Cohen’s *d* = 0.85; 82.2% individually significant; Figure 4). All 11 tissues show positive mean transfer (tissue-level bootstrap 95% CI [0.089, 0.189]). The confidence intervals account for dependence among drug-tissue pairs by resampling at the drug and tissue level rather than treating pairs as independent observations.

**Figure 4:**
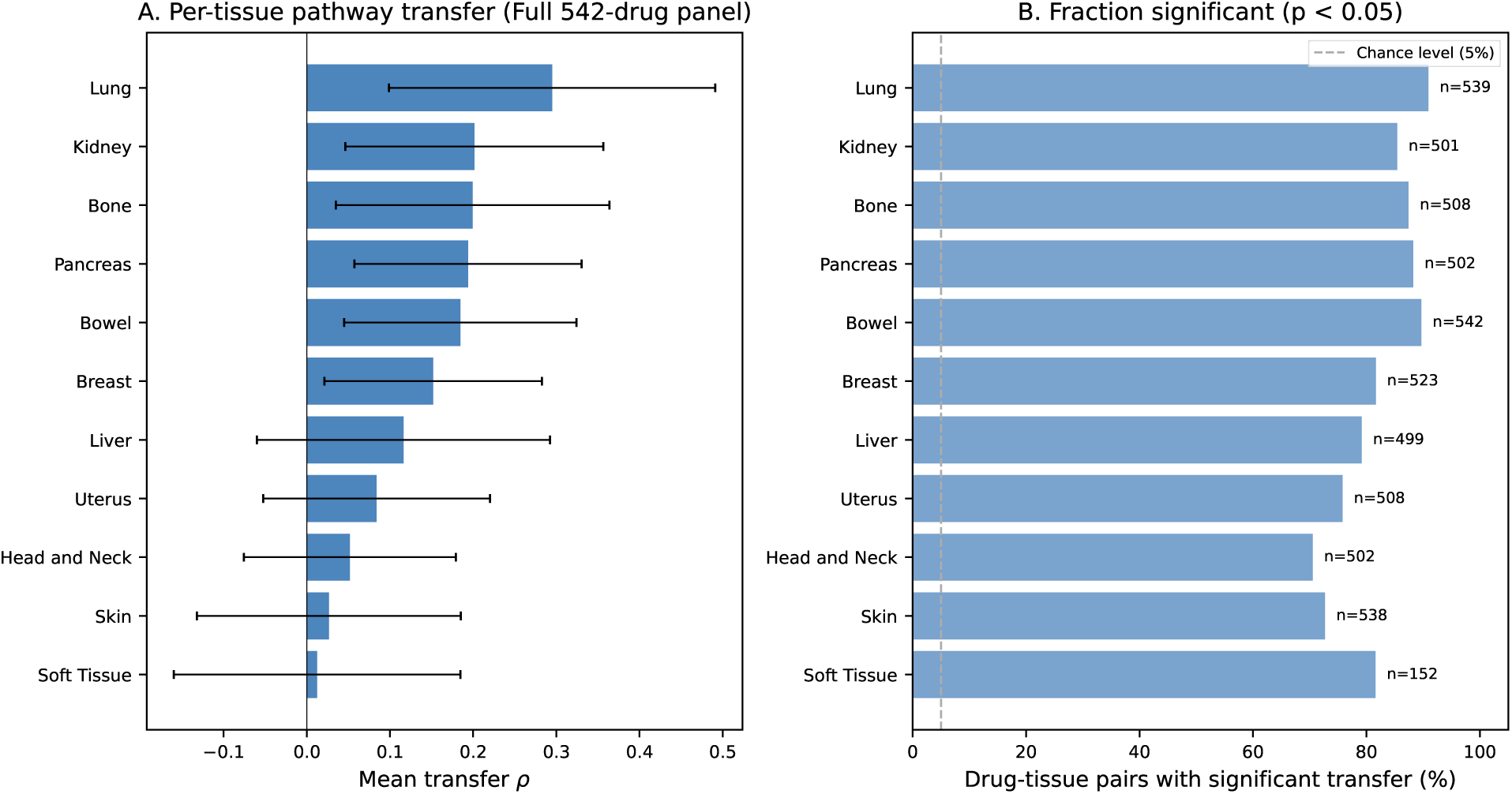
Leave-tissue-out pathway transfer across 542 drugs and 11 tissues (5,314 drug-tissue pairs). (A) Mean transfer *ρ* by held-out tissue. (B) Fraction of drug-tissue pairs with significant positive transfer (*p <* 0.05).

### 3.3 Predictive Performance

Table 3 presents a benchmark following DrEval-style evaluation [Hostallero et al., 2024]: all ML baselines include drug identity as a feature.

**16-drug benchmark.** XGBoost achieves the lowest RMSE (1.185 ± 0.012). GOPA Euclidean (1.226 ± 0.007, 10 seeds) is the strongest neural model. No neural architecture exceeds XGBoost. These results generalize the DrEval finding: tree-baseline superiority is a property of the pharmacogenomic prediction task under fair evaluation.

**Matched pathway baselines.** To isolate representation quality from learner capacity, we evaluate seven pathway representations with matched downstream learners (Table 2). With XGBoost, simple gene-set projections (column-normalized *x* · *B*, mean GO expression) achieve RMSE 1.246, compared to GOPA softmax attention at 1.274. With Ridge, softmax attention achieves a modest edge over shuffled controls (RMSE 1.451 vs. 1.464; residual Pearson 0.233 vs. 0.188). These results establish that GOPA’s predictive advantage stems from its end-to-end architecture (learned drug embeddings, cell encoder, gated aggregation), not from representational superiority of the softmax attention over simpler pathway scores.

**Figure 5:**
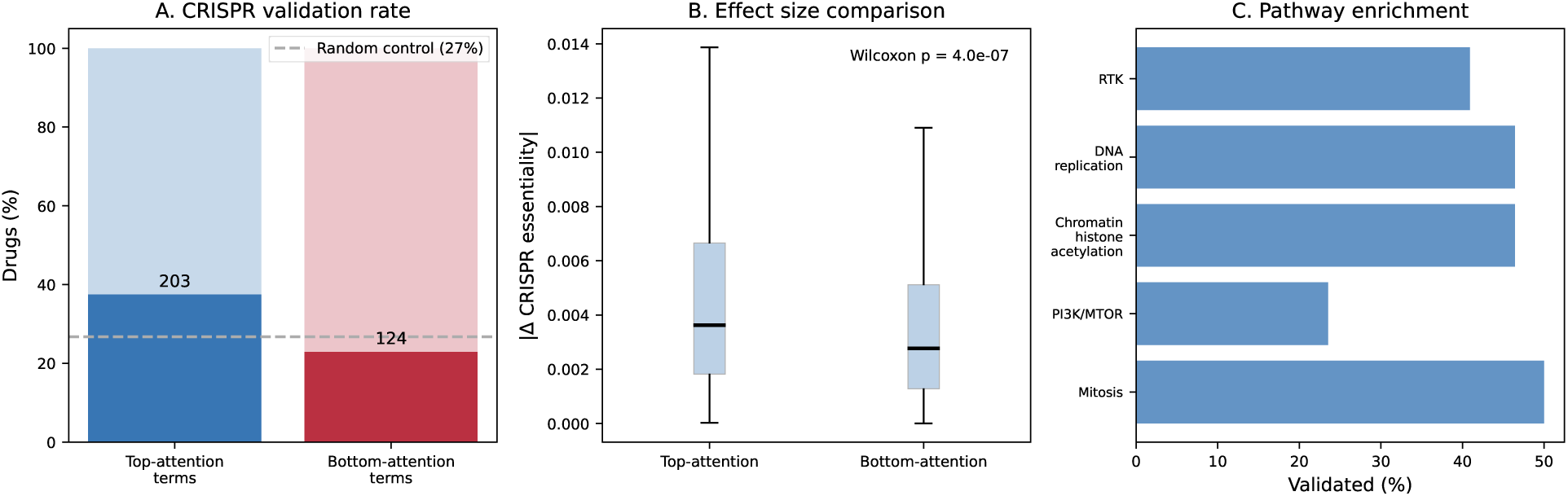
CRISPR dependency validation across 542 drugs. (A) Top-attention pathways show higher CRISPR validation rates (37.5%; 95% CI [33.4, 41.5]) than bottom-attention controls (22.9%; [19.4, 26.4]). (B) Top-attention terms have 1.57× larger |Δ_CRISPR_| than random GO term baselines (*p* = 1.5 × 10^−19^).

**Full 542-drug panel.** On the full GDSC panel (Table 4; 375,885 response rows), XGBoost with one-hot drug encoding (1.713) collapses below the mean-per-drug baseline (1.443), while target-encoded XGBoost (1.322) matches GOPA (1.327). GOPA achieves higher residual Pearson (0.469; 95% CI [0.453, 0.485]) than target-encoded XGBoost (0.391; [0.379, 0.404]); paired difference +0.078 [0.062, 0.093]; GOPA wins on 363 of 539 drugs (67.3%); Wilcoxon *p* = 1.1 × 10^−23^.

### 3.4 Architecture and Geometry

The architecture ablation (Table 6) isolates each component’s contribution. Removing the Gene-Ontology Bridge produces the largest degradation (ΔRMSE = +0.011). Removing the GO DAG GNN (−0.002) or per-drug bias (−0.003) is within seed variance.

Extended branch-dependence experiments (Table 5) separate structural interpretability from predictive dependence. Replacing softmax attention with uniform 1*/T* weights changes RMSE by only +0.001, and permuting attention across samples (+0.004) or shuffling the bridge output (+0.004) has similarly negligible effect. The direct cell-expression branch supplies most of the cell-line-specific predictive signal: removing it while retaining drug and pathway inputs (drug + pathway, no cell) raises RMSE by +0.206. Removing both drug identity and pathway inputs (cell only) collapses prediction entirely (RMSE 2.609; Pearson *r* = 0.197), primarily because drug identity is lost. Cell-specific softmax selectivity contributes negligibly (ΔRMSE ≤ 0.004). GOPA’s pathway value is therefore stable population-level biological summarization rather than prediction routing.

Per-seed matched comparisons (Table 7, 10 seeds) show Euclidean and hyperbolic geometries achieve equivalent prediction (ΔRMSE = +0.008; 95% CI [−0.004, +0.020]; *p* = 0.18). Hyperbolic embeddings preserve GO graph distances better than Euclidean (Table 8, Figure 7): graph-distance correlation *ρ* = 0.732 ± 0.067 vs. 0.474 ± 0.008, edge reconstruction AUROC 0.729 vs. 0.711. However, Euclidean embeddings achieve stronger ancestor retrieval (MRR 0.142 vs. 0.131; P@5 0.128 vs. 0.114) and depth-radius correlation (+0.055 vs. +0.007). Negative controls (randomized GO graph, shuffled annotations) show near-zero hierarchy signal, confirming both signals derive from real ontology structure.

**Table 2:**
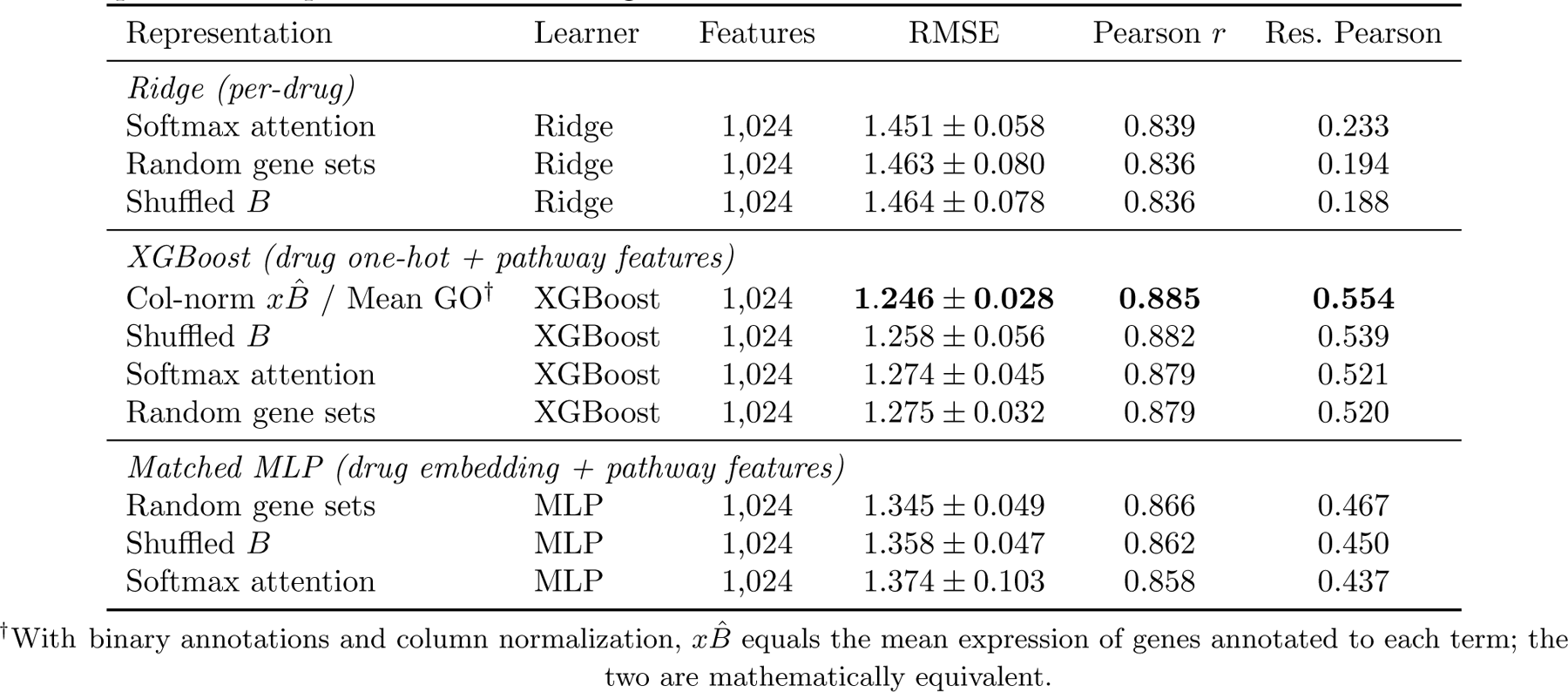
Matched pathway baseline comparison (16-drug benchmark, leave-cell-line-out, 3 seeds). Each representation is evaluated with three downstream learners to isolate representation quality from learner capacity. With XGBoost, simple gene-set projections achieve equivalent or better prediction than GOPA’s softmax attention, indicating the softmax computation does not provide a predictive representation advantage.

**Figure 6:**
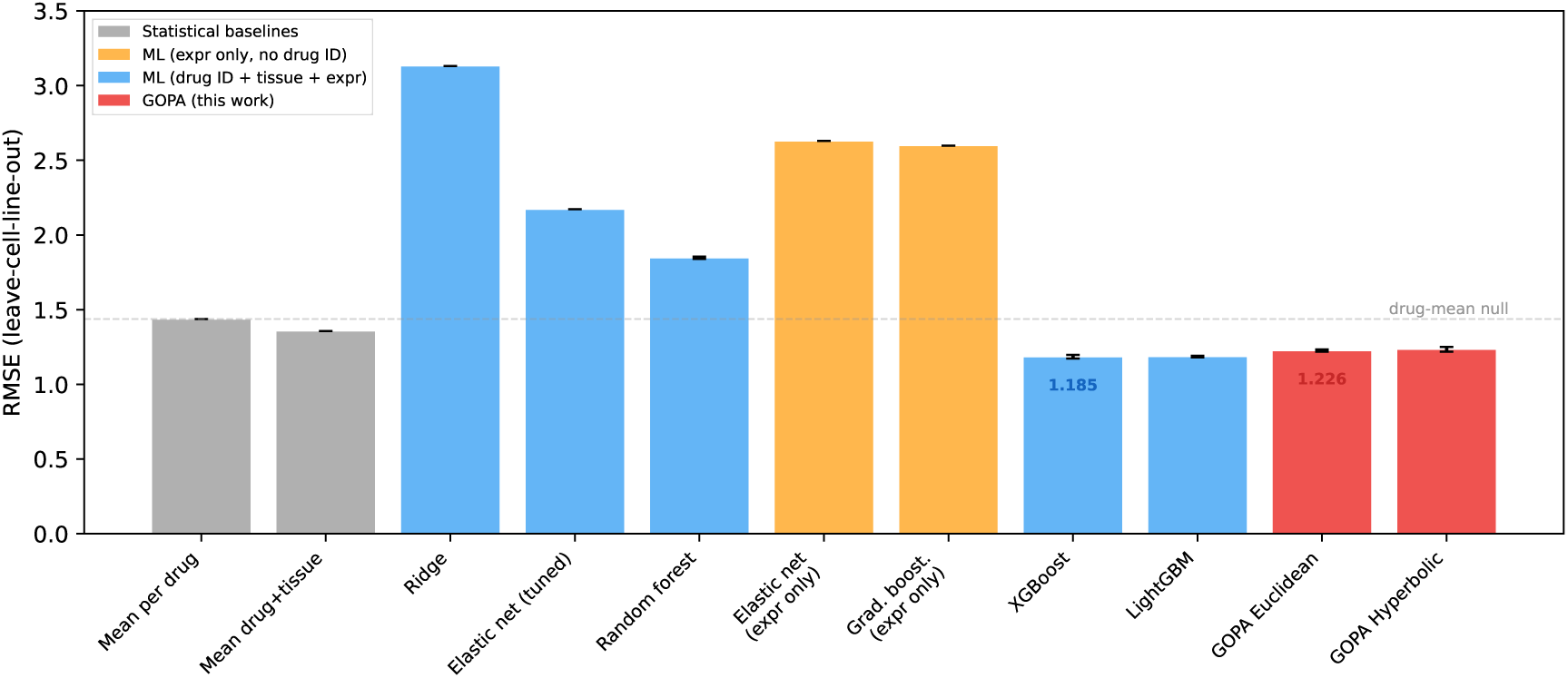
16-drug benchmark RMSE comparison. XGBoost with drug one-hot encoding achieves the lowest RMSE (1.185); GOPA Euclidean (1.226) is the strongest neural model. Full 542-drug panel results including target-encoded XGBoost and paired residual Pearson comparisons are in Table 4.

**Table 3:** Comprehensive DrEval-style benchmark (leave-cell-line-out, 16-drug panel). All ML baselines include drug identity as a feature except where noted, following the principle that properly configured baselines must control for drug-level confounds [Hostallero et al., 2024]. “Res. Pearson” measures cell-line-specific prediction quality after subtracting per-drug mean response (0.000 for the drug-mean null). GOPA results: mean ± s.d. over 10 seeds; baselines over 3 seeds. Best overall RMSE in bold; best GOPA RMSE underlined.

| Model | Category | RMSE | Pearson | Res. Pearson |
| --- | --- | --- | --- | --- |
| *Non-ML baselines* |  |  |  |  |
| Mean per drug | Statistical | 1.437 | 0.841 | 0.000 |
| Mean drug + tissue | Statistical | 1.357 | 0.859 | 0.328 |
| Mean drug + cell line | Statistical | 1.437 | 0.841 | 0.005 |
| *ML baselines (expression only, no drug identity)* |  |  |  |  |
| Elastic net | Linear | 2.629 | 0.170 | 0.121 |
| Gradient boosting | Ensemble | 2.598 | 0.202 | – |
| *ML baselines (drug identity + tissue + expression)* |  |  |  |  |
| XGBoost | Ensemble | **1.185 ± 0.012** | 0.895 | 0.565 |
| LightGBM | Ensemble | 1.186 ± 0.005 | 0.895 | 0.565 |
| Random forest | Ensemble | 1.847 ± 0.001 | 0.863 | 0.149 |
| Elastic net (tuned) | Linear | 2.172 | 0.690 | 0.121 |
| Ridge (tuned) | Linear | 3.131 | 0.547 | 0.085 |
| *GO-pathway attention models (this work)* |  |  |  |  |
| GOPA Euclidean | GO DAG GNN | <u>1.226 ± 0.007</u> | 0.889 ± 0.002 | 0.524 ± 0.012 |
| GOPA Hyperbolic | GO DAG GNN | 1.234 ± 0.016 | 0.887 ± 0.003 | 0.523 ± 0.004 |

This mixed result reflects the GO DAG’s structure: it is a DAG with multiple parents and shortcut edges, not a pure tree. Hyperbolic geometry excels at global distance preservation, while the multiple-parent structure may favor Euclidean space for local ancestor relationships. The practical design guideline: use Euclidean for prediction; use hyperbolic when faithful global distance preservation is the objective.

### 3.5 SHAP Comparison

A natural question is whether post-hoc interpretation of tree models could recover the same biological signals as GOPA’s built-in attention. We train per-drug XGBoost models [Chen and Guestrin, 2016] on expression features, compute SHAP importance per gene [Lundberg and Lee, 2017], and project to GO terms via the annotation matrix. SHAP-GO and GOPA attention show low agreement (mean Spearman *ρ* = 0.115 ± 0.065 across 539 drugs), confirming they produce different pathway rankings. CRISPR functional support rates are comparable (SHAP 22%, GOPA 19%), suggesting both capture biology through different routes. The low agreement does not establish which method is more correct; rather, it indicates that the deterministic projection (softmax(*x_c_* ·*B̂*)) and post-hoc feature attribution provide complementary pathway views. GOPA’s structural advantage is population comparability: the same attention weights apply to any cell line without retraining, whereas SHAP attributions depend on the trained model, hyperparameters, and back-ground distribution (Supplement S25).

**Table 4:** Full GDSC panel results (542 drugs, 702 cell lines, 375,885 response rows, leave-cell-line-out). Residual Pearson measures cell-line-specific prediction after removing per-drug mean response. GOPA results: mean ± s.d. over 3 seeds. Paired comparison uses drug-level bootstrap.

| Model | RMSE | Pearson | Res. Pearson | 95% CI |
| --- | --- | --- | --- | --- |
| Mean per drug | 1.443 | 0.842 | — | — |
| XGBoost (one-hot drug) | $1.713 \pm 0.009$ | 0.802 | 0.151 | — |
| Target-enc. XGBoost | $1.322 \pm 0.012$ | 0.870 | 0.391 | [0.379, 0.404] |
| Euclidean GOPA | $1.327 \pm 0.002$ | 0.869 | 0.469 | [0.453, 0.485] |
| Hyperbolic GOPA | $1.326 \pm 0.009$ | 0.868 | 0.450 | — |
| *Paired comparison (Euclidean GOPA vs. target-encoded XGBoost)* |  |  |  |  |
| Difference: +0.078 [0.062, 0.093]; GOPA wins 363/539 drugs; Wilcoxon $p = 1.1 \times 10^{-23}$ | | | | |

**Table 5:**
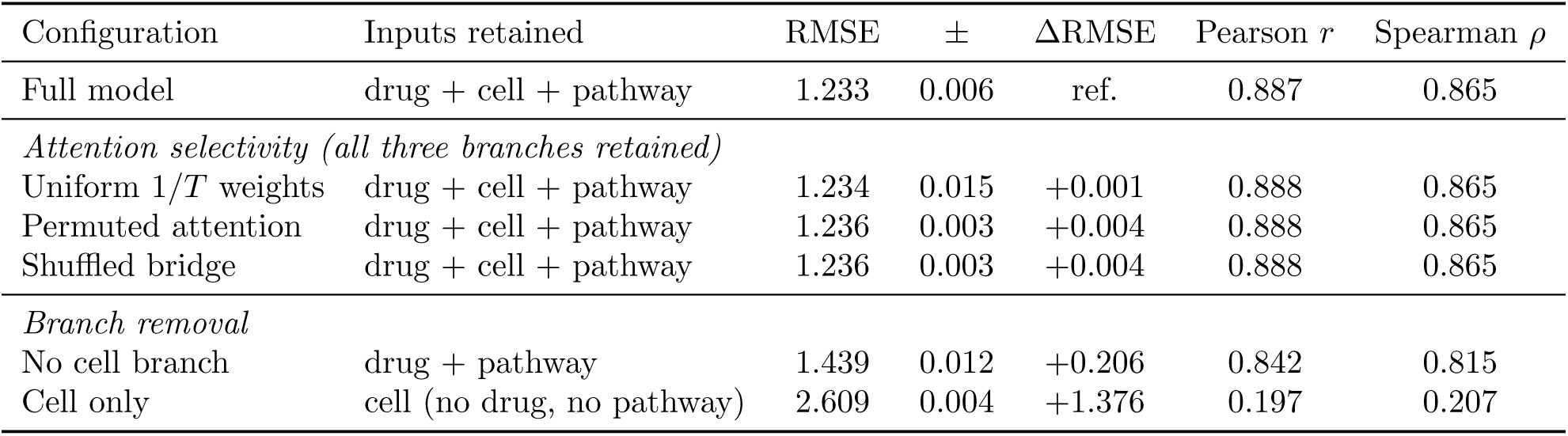
Branch-dependence ablations (16-drug benchmark, 30 epochs, 3 seeds). Each row states the inputs retained. Attention selectivity contributes negligibly; the direct cell branch supplies most cell-line-specific signal.

| Configuration | Inputs retained | RMSE | ± | ΔRMSE | Pearson $r$ | Spearman $\rho$ |
| --- | --- | --- | --- | --- | --- | --- |
| Full model | drug + cell + pathway | 1.233 | 0.006 | ref. | 0.887 | 0.865 |
| *Attention selectivity (all three branches retained)* |  |  |  |  |  |  |
| Uniform $1/T$ weights | drug + cell + pathway | 1.234 | 0.015 | +0.001 | 0.888 | 0.865 |
| Permuted attention | drug + cell + pathway | 1.236 | 0.003 | +0.004 | 0.888 | 0.865 |
| Shuffled bridge | drug + cell + pathway | 1.236 | 0.003 | +0.004 | 0.888 | 0.865 |
| *Branch removal* |  |  |  |  |  |  |
| No cell branch | drug + pathway | 1.439 | 0.012 | +0.206 | 0.842 | 0.815 |
| Cell only | cell (no drug, no pathway) | 2.609 | 0.004 | +1.376 | 0.197 | 0.207 |

### 3.6 Additional Validation

**Molecular drug encoding.** Replacing learned embeddings with Morgan fingerprints (1024-bit, radius 2) matches leave-cell-line-out performance (1.237 vs. 1.226) and improves leave-drug-out generalization (1.991 vs. 2.277; Supplement S7).

**External validation.** Cross-database validation (GDSC1→GDSC2) retains strong ranking for overlapping drugs (Pearson *r* = 0.817, 122 drugs) but near-chance for unseen drugs (*r* = 0.16; Supplement S6).

**gCSI locked external validation.** To test the defense-response hypothesis on an independent pharmacogenomic panel, we evaluated gCSI [Haverty et al., 2016] (Genentech Cell Line Screening Initiative; 268 cell lines, 7 overlapping kinase drugs). The hypothesis (defense activity predicts kinase-inhibitor sensitivity) and direction were locked from GDSC before inspecting gCSI results. The gCSI AUC convention was empirically verified as activity area (higher AUC = greater sensitivity; AUC-IC50 Spearman *ρ* = −0.777; A375 BRAF-mutant AUC = 0.84 for MEK inhibitor).

**Table 6:**
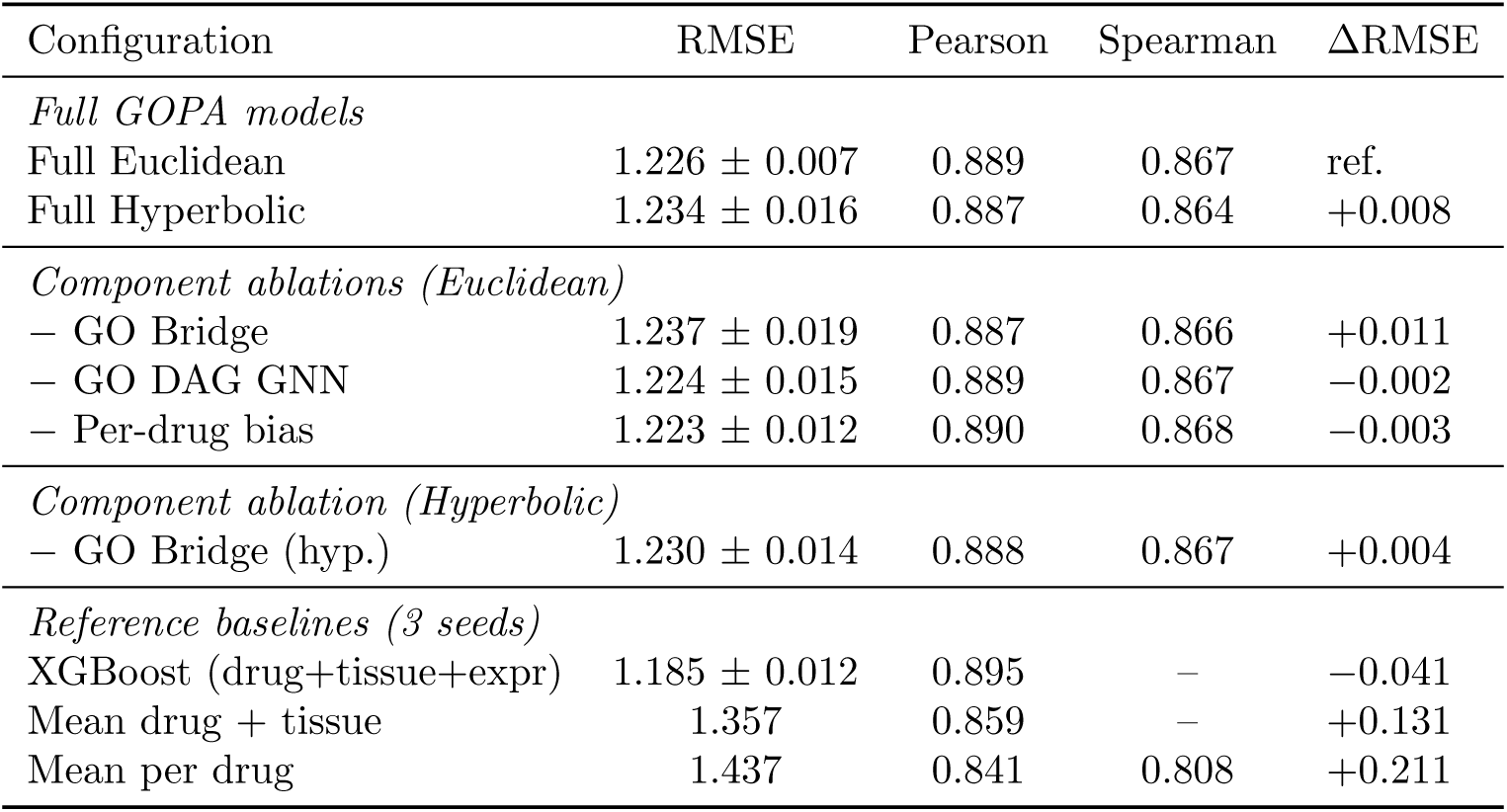
Architecture ablation (leave-cell-line-out, 16-drug benchmark). Each row removes one component from the full GOPA model. Removing the GO Bridge causes the largest degradation within the GOPA family. Mean ± s.d. over 10 seeds.

**The locked gCSI test did not reproduce the consistent association observed in GDSC and PRISM**: mean defense-response AUC *ρ* = −0.107 (5 of 7 kinase drugs showed negative AUC correlation, indicating defense-resistance association under the verified convention; 2 of 7 showed positive AUC correlation, consistent with the GDSC sensitivity direction). Per-drug results: Crizotinib *ρ* = −0.167 (*p* = 0.006), Erlotinib −0.089 (ns), GDC-0941 +0.062 (ns), Lapatinib −0.217 (*p <* 0.001), PD-0325901 −0.085 (ns), Rapamycin +0.008 (ns), BI-D1870 −0.260 (*p <* 0.001). A methodological control confirmed this is not a scoring artifact: the same sum-of-Z-scores method applied to GDSC recovers the expected negative LN IC50 correlations for all overlapping drugs (Crizotinib *ρ* = −0.227, Lapatinib −0.126). The gCSI panel lacks the MEK inhibitors (Trametinib, Refametinib, Selumetinib) that showed the strongest GDSC confound-resistant signals, uses a different response metric (AUC vs. LN IC50), and contains 268 cell lines (vs. *>*900 in GDSC). The non-reproduction is consistent with assay, panel-composition, response-metric, or drug-set dependence.

**Multiomic integration.** We evaluate four modality configurations on the full GDSC panel: expression only, expression + copy-number, expression + mutation, and expression + copy-number + mutation. Expression alone provides the best performance (RMSE 1.327 ± 0.002). Adding gene-level copy-number ratios degrades accuracy modestly (1.340 ± 0.005), as does adding binary somatic mutation indicators (1.338 ± 0.004). The combined three-modality configuration is weakest (1.345 ± 0.006). This pattern likely reflects two factors: (1) gene dosage effects are already captured in log-TPM expression, making copy-number features redundant; (2) binary mutation indicators are extremely sparse (mean *<* 3% of genes mutated per cell line), contributing noise relative to the 512-dimensional expression input. The GO annotation matrix *B* is defined over expression genes only; copy-number and mutation features receive zero GO annotation weights, diluting the pathway signal. Future work could explore pathway-aggregated copy-number features or mutation burden scores that align with the GO term structure (Supplement S5).

**Table 7:**
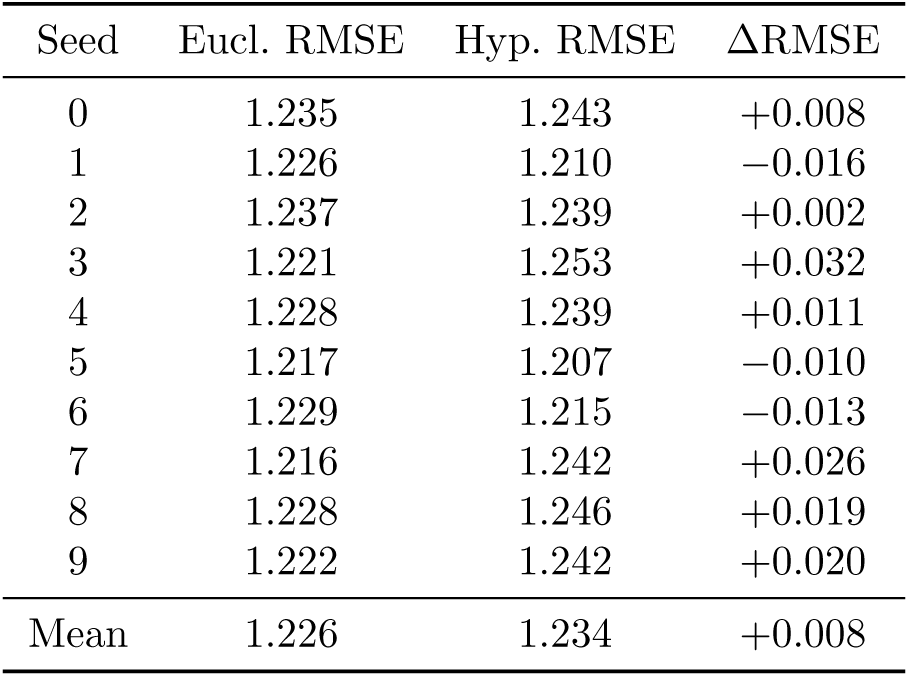
Matched hyperbolic versus Euclidean per-seed deltas (leave-cell-line-out, 10 seeds). Positive ΔRMSE means hyperbolic is worse. Prediction performance is comparable (*p* = 0.18, paired *t*-test); the hyperbolic advantage is in representation quality (Table 8).

**Table 8:**
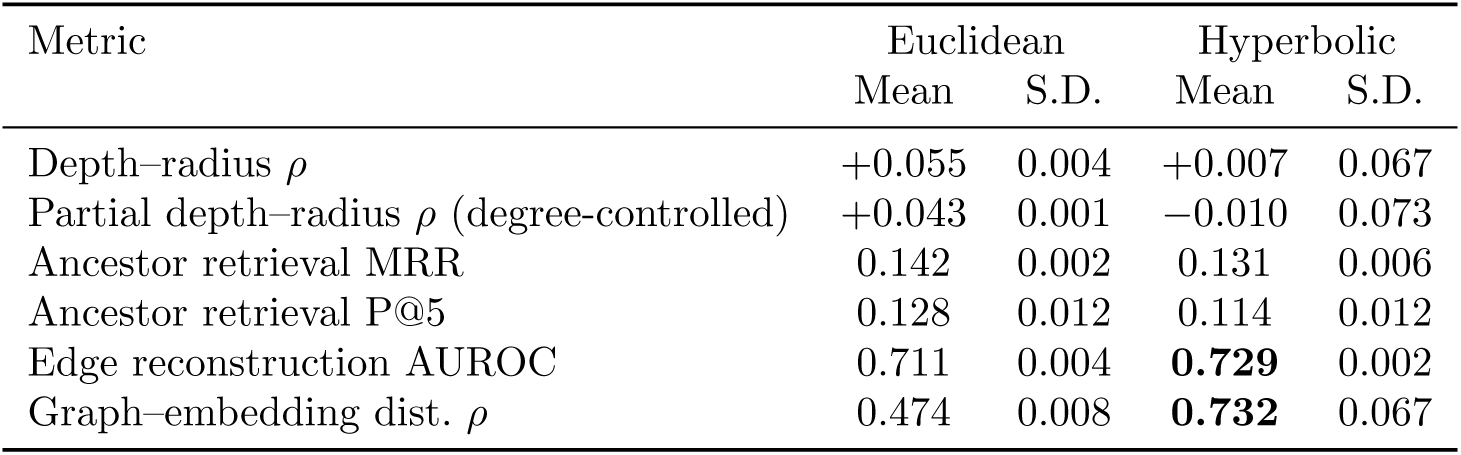
Comprehensive hierarchy audit of GO term embeddings (mean ± s.d. over 3 seeds). Graph-distance correlation (Spearman between GO DAG shortest-path distance and embedding distance) is the strongest hierarchy metric. Hyperbolic embeddings preserve graph distances 50% better than Euclidean (*ρ* = 0.732 vs. 0.474) and achieve higher edge reconstruction AUROC. Degree-confound-controlled d<u>epth–radius partial correlation confir</u>m<u>s the signal is not driven by node d</u>egree.

**Computational cost.** Table 9 summarizes computational costs across components and scales. Deterministic pathway activity computation processes 11,248 samples with 512 GO terms in 0.07 seconds on CPU (165,000 samples/s), scaling linearly in both samples and terms. GOPA inference runs at 0.002 seconds per batch of 256 samples (∼242K parameters). Full training on the 16-drug benchmark takes ∼25 seconds per seed on CPU; the 542-drug panel takes ∼14 minutes. For comparison, Ridge training takes 0.04 seconds and XGBoost (500 trees) takes ∼8 seconds.

**Figure 7:**
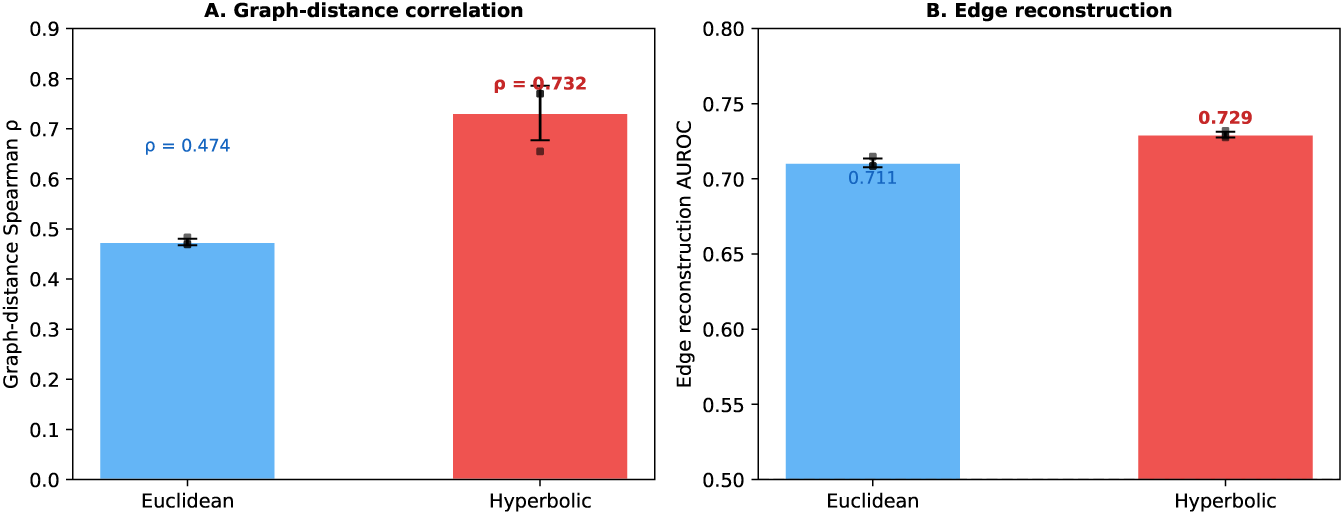
GO term embedding geometry. Hyperbolic embeddings preserve global graph distances better than Euclidean (*ρ* = 0.732 vs. 0.474) and achieve higher edge reconstruction AUROC (0.729 vs. 0.711), while Euclidean embeddings show stronger ancestor retrieval (MRR 0.142 vs. 0.131) and depth-radius correlation (+0.055 vs. +0.007). Negative controls confirm both signals derive from real ontology structure.

**Table 9:**
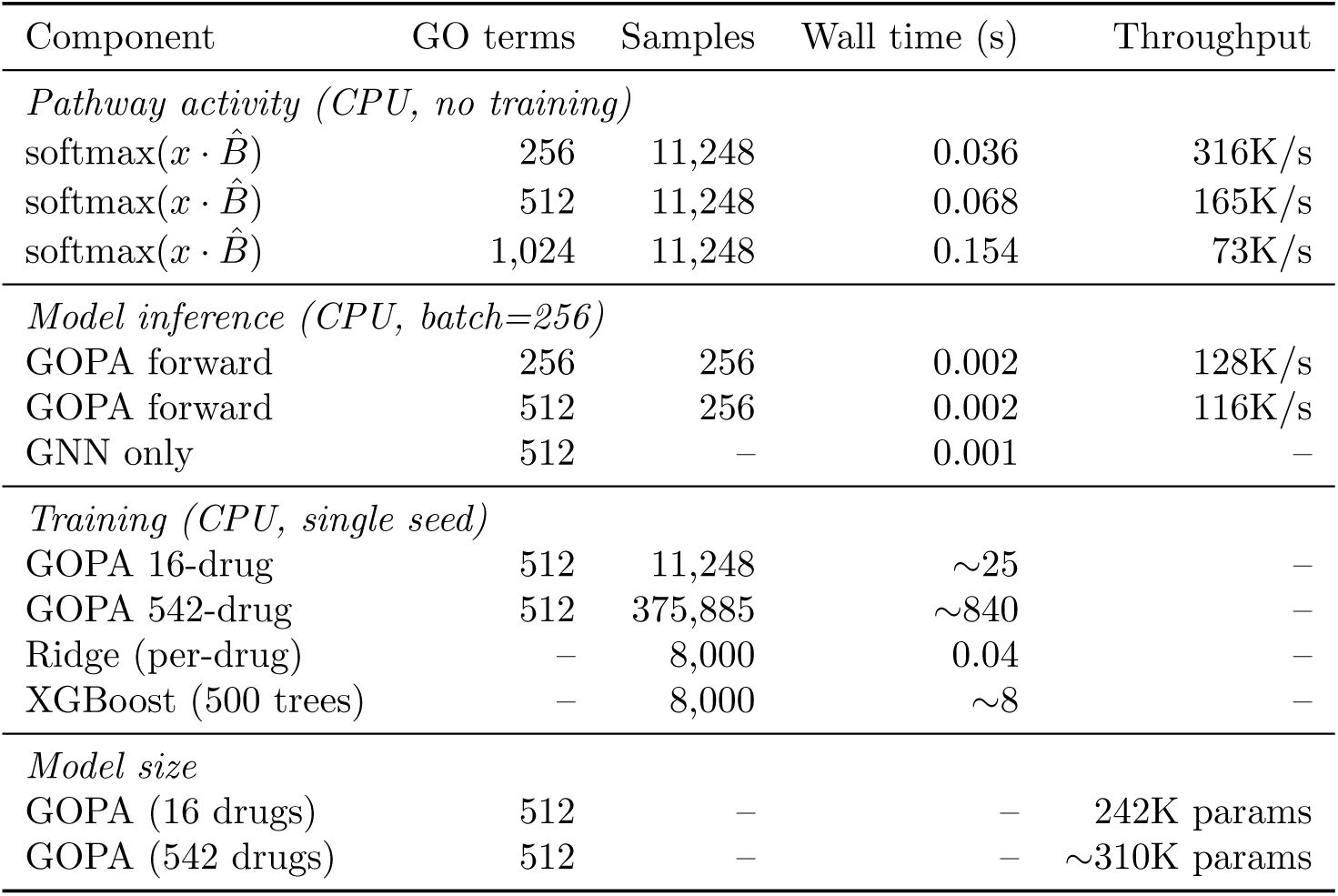
Computational cost summary. Pathway activity is deterministic (no GPU required). All timings on a single CPU (Intel Xeon).

| Component | GO terms | Samples | Wall time (s) | Throughput |
| --- | --- | --- | --- | --- |
| *Pathway activity (CPU, no training)* |  |  |  |  |
| $\text{softmax}(x \cdot \hat{B})$ | 256 | 11,248 | 0.036 | 316K/s |
| $\text{softmax}(x \cdot \hat{B})$ | 512 | 11,248 | 0.068 | 165K/s |
| $\text{softmax}(x \cdot \hat{B})$ | 1,024 | 11,248 | 0.154 | 73K/s |
| *Model inference (CPU, batch=256)* |  |  |  |  |
| GOPA forward | 256 | 256 | 0.002 | 128K/s |
| GOPA forward | 512 | 256 | 0.002 | 116K/s |
| GNN only | 512 | — | 0.001 | — |
| *Training (CPU, single seed)* |  |  |  |  |
| GOPA 16-drug | 512 | 11,248 | ~25 | — |
| GOPA 542-drug | 512 | 375,885 | ~840 | — |
| Ridge (per-drug) | — | 8,000 | 0.04 | — |
| XGBoost (500 trees) | — | 8,000 | ~8 | — |
| *Model size* |  |  |  |  |
| GOPA (16 drugs) | 512 | — | — | 242K params |
| GOPA (542 drugs) | 512 | — | — | ~310K params |

## 4 Discussion

### 4.1 Defense Response and MEK Inhibitor Sensitivity

The association between defense response pathways and kinase inhibitor sensitivity, observed across 12 drugs in GDSC and supported by PRISM cross-assay transfer (11 of 11 overlapping drugs), is the study’s primary biological finding. The effect is strongest and most confound-resistant for MEK inhibitors (Refametinib partial *ρ* = −0.204, Trametinib −0.181 after five-confound correction), consistent with the MAPK/PD-L1 model [Stutvoet et al., 2019]: MAPK/ERK signaling is associated with PD-L1 expression, and cell lines with high defense response expression show greater sensitivity to MEK inhibition. The 72% signal retention after controlling for IFN/JAK-STAT, proliferation, HLA, EMT, and CD274 simultaneously indicates that the association is not fully explained by the five measured confounds.

However, a locked test on gCSI (268 cell lines, 7 kinase drugs, AUC response metric) did not reproduce the consistent association (Section 3.6), limiting the generalizability claim. The CD274 analysis shows an association consistent with MAPK biology but does not constitute mechanistic confirmation; CRISPR knockout of defense response genes during MEK inhibitor treatment would provide direct perturbation evidence.

### 4.2 What Deterministic Pathway Projection Contributes

GOPA’s parameter-free attention (softmax(*x_c_* · *B̂*)) provides structurally grounded interpretive properties that learned attention mechanisms and post-hoc methods do not share: the weights are computable without model inference, invariant to training, and population-comparable. GOPA and SHAP-projected pathway rankings show low agreement (*ρ* = 0.115), with comparable CRISPR functional support rates (SHAP 22%, GOPA 19%), suggesting complementary rather than competing biological views.

Matched pathway baseline experiments establish that GOPA’s softmax attention does not provide a predictive representation advantage over simpler projections (column-normalized *x* · *B*, mean GO expression) when evaluated with the same downstream learner. With XGBoost, all projections achieve equivalent RMSE (∼1.246 vs. 1.274 for softmax attention). Branch-dependence ablations reinforce this: replacing softmax with uniform 1*/T* weights changes RMSE by only +0.001, establishing that the bridge provides a population-level pathway representation rather than cell-specific routing. GOPA’s predictive contribution stems from its end-to-end architecture integrating pathway, cell, and drug representations, not from attention selectivity. The softmax concentration provides a modest edge for linear learners (Ridge), enables the defense-response discovery with stronger kinase-specific discrimination than simple scores, and the key value of deterministic, population-comparable pathway summaries.

### 4.3 What Hyperbolic Geometry Contributes

The geometry comparison provides a nuanced answer: Poincaŕe-ball embeddings preserve global GO graph distances substantially better than Euclidean (*ρ* = 0.732 vs. 0.474) and improve edge reconstruction AUROC (0.729 vs. 0.711). However, Euclidean embeddings achieve stronger ancestor retrieval (MRR 0.142 vs. 0.131; P@5 0.128 vs. 0.114) and depth-radius correlation (+0.055 vs. +0.007). Prediction is statistically indistinguishable (ΔRMSE = +0.008; *p* = 0.18), confirming the predictive signal flows through the annotation-based attention mechanism, which is geometry-agnostic. The GNN-refined embeddings enter only through gated aggregation downstream and contribute negligibly to prediction (ΔRMSE = −0.002). This finding contributes to the growing evidence that geometric fidelity and task performance can decouple when the task-relevant computation does not pass through the geometry-sensitive component.

### 4.4 Benchmarking and the Drug-Encoding Confound

Properly configured baselines with drug identity features substantially outperform expression-only baselines (XGBoost 1.185 vs. elastic net 2.629 on 16 drugs), confirming DrEval principles [Hostallero et al., 2024]. On the full panel, target-encoded XGBoost matches GOPA on RMSE (1.322 vs. 1.327); the one-hot encoding, not the tree architecture, was the bottleneck. GOPA achieves higher residual Pearson (0.47 vs. 0.39; paired difference +0.08 [0.06, 0.09]; GOPA wins on 67% of drugs; Wilcoxon *p <* 10^−23^), indicating statistically significant cell-line-specific prediction advantage. Tree models with thoughtful drug encoding are competitive on aggregate RMSE, but GOPA provides interpretable pathway attention that tree models lack.

### 4.5 Study Limitations

On the 16-drug benchmark, GOPA does not exceed tree baselines on RMSE, and on 542 drugs it matches target-encoded XGBoost. Matched pathway baseline experiments show that simple gene-set projections achieve equivalent prediction with the same downstream learner, limiting GOPA’s novelty claim to its architectural integration and interpretive properties. The parameter-free design ensures interpretive stability but limits expressiveness; learned attention could capture nonlinear pathway interactions. Expression dominates; multi-omic integration does not improve performance. The defense response discovery, though supported across 12 drugs in GDSC and PRISM and 72% confound-resistant, was not reproduced in gCSI (non-replication with important caveats including different response metric, smaller sample, and older data version), requiring caution about generalizability. Experimental perturbation evidence (CRISPR under drug treatment) remains needed for mechanistic claims. Leave-tissue-out transfer demonstrates cross-tissue generalization but does not substitute for patient-derived validation. PRISM shares a substantial cell-line ecosystem with GDSC; the support represents cross-assay pharmacological evidence, not independent cohort validation. The “cell-intrinsic inflammatory” framing reflects the cell-line context; whether this corresponds to tumor-intrinsic programs in patient tumors, where immune infiltration confounds the measurement, remains to be established. The mixed geometry result (hyperbolic better on global distances, Euclidean better on ancestor retrieval) means neither geometry is uniformly superior for representing GO structure.

## 5 Methods

### 5.1 Data and Harmonization

We use DepMap expression profiles (TPM, log-transformed), GDSC release 8.5 fitted dose-response (LN IC50), and GO Basic ontology with GOA human annotations. Gene-term annotations are propagated along the GO DAG following the true path rule: a gene annotated to term *T* also receives annotations for all ancestors of *T* within the selected term set. For multi-omic experiments, we additionally use gene-level copy-number ratios and somatic mutation calls (binary). Drug target pathway metadata is from the GDSC screened compounds manifest. Cell-line matching via normalized name matching retains 466 cell lines for the 16-drug benchmark (45.7% of DepMap models). The full GDSC panel retains 702 cell lines across 542 drugs (Supplement S1-S2).

**Table 10:**
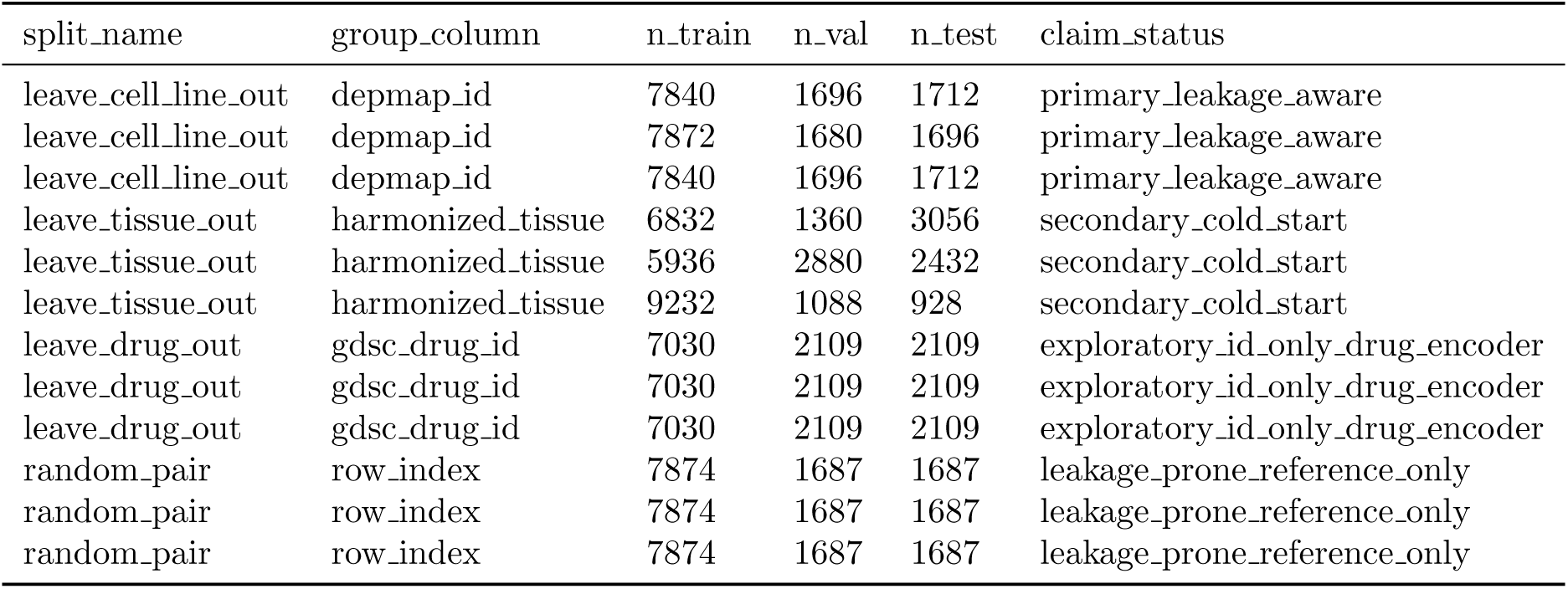
Materialized split design and claim status.

### 5.2 Benchmark Design

**Selected-drug benchmark.** 11,248 response rows across 16 drugs and 512 expression features.

**Full GDSC panel.** All GDSC drugs with ≥50 cell lines per drug: 542 drugs, 702 cell lines, 375,885 response rows.

**Split design.** Leave-cell-line-out splitting assigns all rows per cell line exclusively to train (70%), validation (15%), or test (15%). Features are standardized using training-fold statistics only. Leave-tissue-out and leave-drug-out splits serve as secondary stress tests.

### 5.3 Architecture

Figure 8 provides an overview of the GOPA architecture. Let 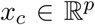, denote expression for cell line *c*, *d* a drug index, 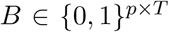 the gene-term annotation matrix with ancestor-propagated annotations (*T* = 512 terms selected by annotation prevalence), 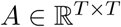 the GO DAG adjacency matrix (symmetric, with self-loops), and *y_cd_*the response value.

**GO term embeddings.** Initial term features (annotation prevalence, ontology depth) are projected to R*^h^* and refined through *L* = 2 layers of GO DAG graph convolution:

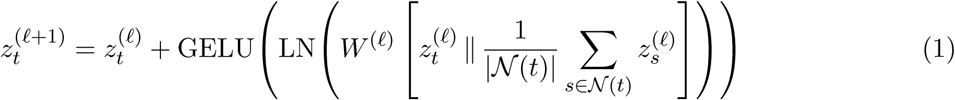

**Gene-Ontology Bridge.** Column-normalized annotation matrix: 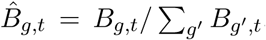. For each sample, pathway activations are 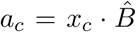, softmax-normalized to attention weights 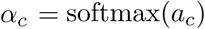, which contain *no learned parameters*. The attention weights aggregate learned GO term embeddings:

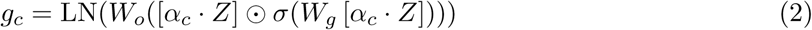

**Prediction.** 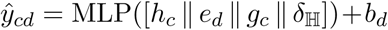, where *h_c_* is the cell encoder output (2-layer MLP), *e_d_* the drug embedding, *b_d_* a per-drug bias, and 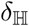 contains pairwise Poincaŕe distances (hyperbolic mode only). Prediction head: 3-layer MLP (3*h* → *h* → *h/*2 → 1) with LayerNorm, GELU, and dropout. Hidden dimension *h* = 128. Total parameters: ∼242K (16 drugs), ∼310K (542 drugs).

**Figure 8:**
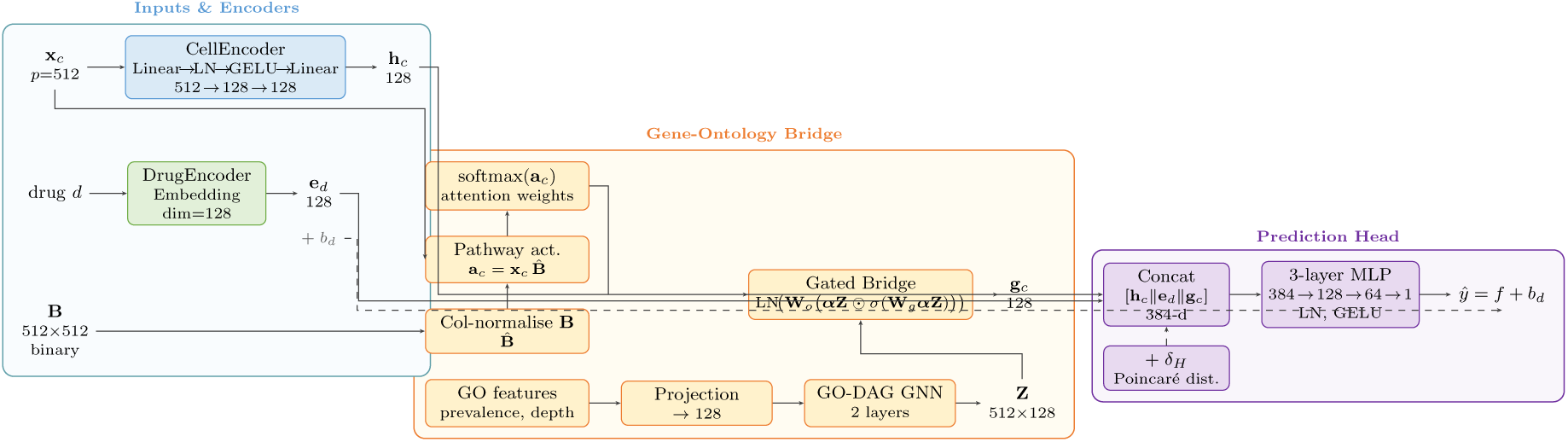
GOPA architecture. Gene expression is projected through a cell encoder and through the column-normalized gene-term annotation matrix into deterministic pathway attention weights (softmax(*x_c_* · *B̂*), no learned parameters). GO term embeddings, refined by a GNN on the GO DAG, are aggregated via gated attention in the Gene-Ontology Bridge. The resulting pathway representation is concatenated with cell and drug embeddings for prediction. The pathway attention is computable without model inference; learned parameters enter only through the GO term embeddings, gated aggregation, and prediction head.

### 5.4 Training

AdamW with cosine-annealing learning rate (5 × 10^−4^ peak, 5-epoch warmup), gradient clipping (norm 1.0), weight decay 10^−4^, batch size 256, and early stopping on validation RMSE with patience 25. Features are standardized per training fold. GOPA experiments use 10 random seeds; baselines use 3 seeds. Single CPU, ∼25s/seed (16 drugs), ∼14 min (542 drugs).

### 5.5 Baselines

XGBoost and LightGBM with drug one-hot + tissue one-hot + 512 standardized expression features (*n*_estimators_ = 500, max depth = 6, lr = 0.05, subsample = 0.8). Target-encoded XGBoost replaces one-hot drug encoding with train-fold mean/std/count encoding of drug ID. Matched pathway baselines evaluate seven representations (raw *xB*, column-normalized *xB*, mean GO expression, z-scored mean, softmax attention, shuffled *B*, random gene sets) with Ridge, XGBoost, and matched MLP using the same splits.

### 5.6 Biological Validation

Pathway attention-response correlations (Spearman) are computed per drug across cell lines. PRISM transfer uses separately computed GDSC and PRISM correlation vectors (158 drugs, 477 cell lines; shared cell-line ecosystem). CRISPR: differential gene essentiality (Mann-Whitney) between drug-sensitive/resistant quartiles for genes in top-20 attention GO terms, benchmarked against 50 random draws; DepMap dependency data measure essentiality without drug treatment. Confound control: partial Spearman correlations via OLS residualization against five scores.

#### Algorithm 1

GOPA Training and Evaluation

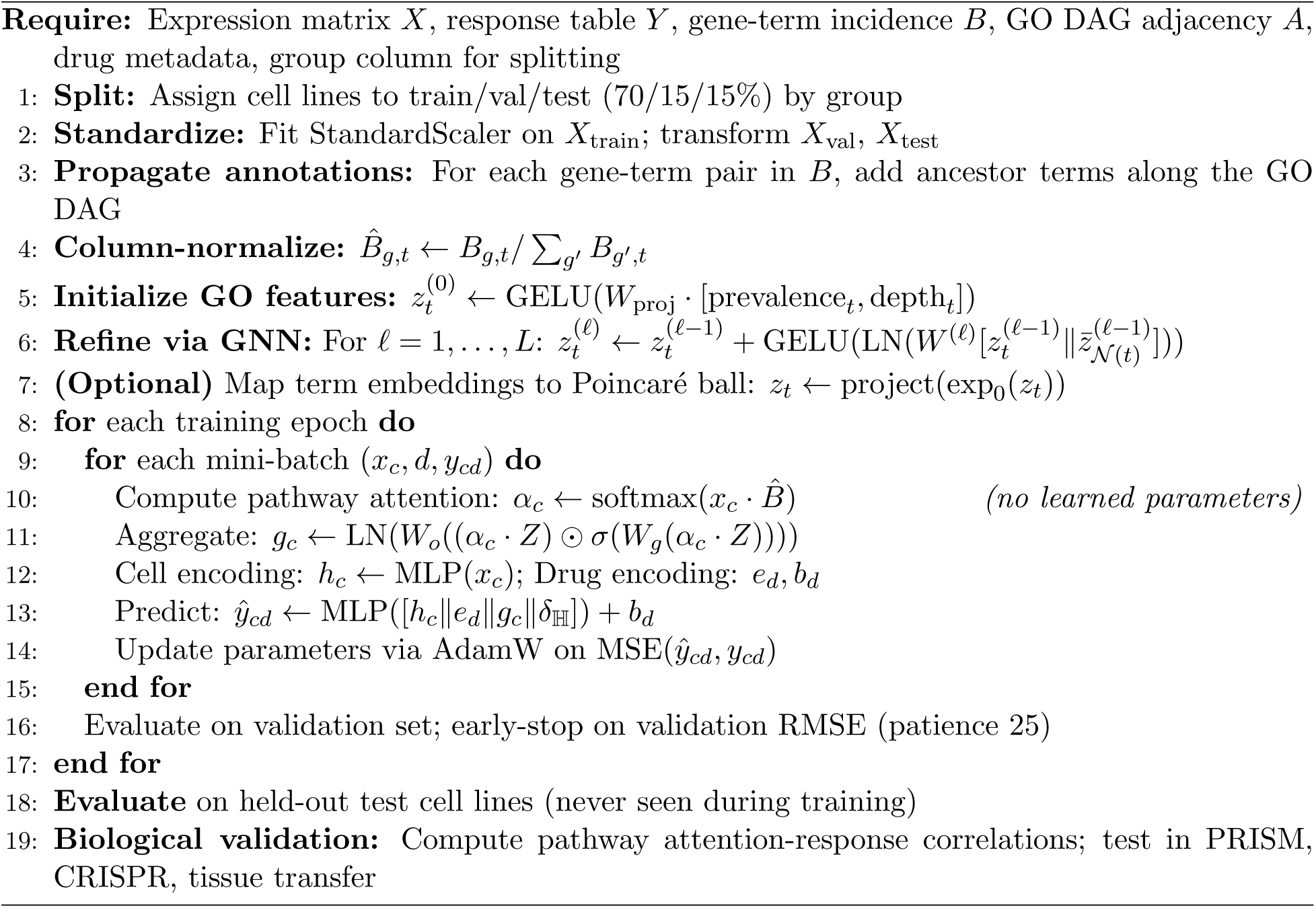

## 6 Conclusion

GOPA provides a deterministic, annotation-guided pathway representation within an end-to-end prediction architecture. The parameter-free attention enables population-scale pathway validation without model retraining. GOPA matches target-encoded XGBoost on full-panel RMSE (1.327 vs. 1.322) with higher residual Pearson (0.47 vs. 0.39; paired difference +0.08 [0.06, 0.09]; Wilcoxon *p <* 10^−23^), though simple gene-set projections achieve equivalent prediction under matched learners, localizing GOPA’s contribution to its architecture and interpretive properties. The defense response association with kinase inhibitor sensitivity, strongest for MEK/MAPK inhibitors (partial *ρ <* −0.15 after five-confound correction), is observed in GDSC and PRISM but was not reproduced in gCSI (non-replication; MEK inhibitors absent from that panel). The finding may therefore be MAPK-pathway-specific and assay-dependent rather than a universal kinase-inhibitor phenomenon. Poincaré-ball embeddings preserve global GO distances better than Euclidean, while Euclidean embeddings achieve stronger ancestor retrieval; neither improves prediction. These results support ontology-guided pathway analysis as a tool for structured biological discovery in pharmacogenomics, while establishing clear boundaries on what the evidence supports: the predictive advantage over simpler representations is architectural rather than representational, the biological discovery is strongest for MEK inhibitors and does not generalize across pharmacogenomic panels, and patient-derived validation remains necessary.

## 7 Data and Code Availability

All author-generated code required to reproduce the analyses, model training, evaluation, statistical tests, tables, and figures is available at https://github.com/ChimdiWalter/GOPA-ontology-pathway-attention The exact version corresponding to this manuscript is tagged as v1.0.0-gopa-preprint. Dataset download and harmonization instructions, fixed split manifests, software dependencies, and reproducibility commands are provided in the repository. Third-party pharmacogenomic datasets (DepMap: https://depmap.org; GDSC: https://www.cancerrxgene.org; GO: http://geneontology.org; gCSI: https://figshare.com/articles/dataset/5297647) are not redistributed and must be obtained from their original sources under the applicable terms.

## Supporting information

Supplementary Material: Tables S1-S9, Figures S1-S2, Sections S1-S29 (split integrity, benchmarks, ablations, biological validation, geometry audit, f

## References

Sara Aibar, Carmen Bravo González-Blas, Thomas Moerman, Vân Anh Huynh-Thu, Hana Imrichova, Gert Hulselmans, Florian Rambow, Jean-Christophe Marine, Pierre Geurts, Jan Aerts, et al. SCENIC: single-cell regulatory network inference and clustering. Nature Methods, 14: 1083–1086, 2017. doi: 10.1038/nmeth.4463.

David A. Barbie, Pablo Tamayo, Jesse S. Boehm, So Young Kim, Susan E. Moody, Ian F. Dunn, Anna C. Schinzel, Peter Sandy, Etienne Meylan, Claudia Scholl, et al. Systematic RNA interference reveals that oncogenic KRAS-driven cancers require TBK1. Nature, 462:108–112, 2009. doi: 10.1038/nature08460.

Jordi Barretina, Giordano Caponigro, Nicolas Stransky, Kavitha Venkatesan, Adam A. Margolin, Sungjoon Kim, Christopher J. Wilson, Joseph Lehar, Gregory V. Kryukov, Dmitriy Sonkin, Anupama Reddy, Manway Liu, Lauren Murray, Michael F. Berger, John E. Monahan, Paula Morais, Jodi Meltzer, Adam Korejwa, Judit Jane-Valbuena, Felipa A. Mapa, Jeremy Thibault, Eva Bric-Furlong, Pichai Raman, Aaron Shipway, Ingo H. Engels, Jill Cheng, Guoying K. Yu, Jianjun Yu, Peter Aspesi, Melanie de Silva, Kalpana Jagtap, Michael D. Jones, Li Wang, Charles Hatton, Emanuele Palescandolo, Supriya Gupta, Scott Mahan, Carrie Sougnez, Robert C. Onofrio, Ted Liefeld, Laura MacConaill, Wendy Winckler, Michael Reich, Nanxin Li, Jill P. Mesirov, Stacey B. Gabriel, Gad Getz, Kristin Ardlie, Vivien Chan, Vic E. Myer, Barbara L. Weber, Jeff Porter, Markus Warmuth, Peter Finan, Jennifer L. Harris, Matthew Meyerson, Todd R. Golub, Michael P. Morrissey, William R. Sellers, Robert Schlegel, and Levi A. Garraway. The cancer cell line encyclopedia enables predictive modelling of anticancer drug sensitivity. Nature, 483: 603–607, 2012. doi: 10.1038/nature11003.

Michael M Bronstein, Joan Bruna, Taco Cohen, and Petar Veličković. Geometric deep learning: Grids, groups, graphs, geodesics, and gauges. arXiv preprint arXiv:2104.13478, 2021.

Ines Chami, Zhitao Ying, Christopher Ré, and Jure Leskovec. Hyperbolic graph convolutional neural networks. In Advances in Neural Information Processing Systems, 2019.

Tianqi Chen and Carlos Guestrin. XGBoost: A scalable tree boosting system. Proceedings of the 22nd ACM SIGKDD, pages 785–794, 2016. doi: 10.1145/2939672.2939785.

Steven M. Corsello, Rohith T. Nagari, Ryan D. Spangler, Jordan Rossen, Mustafa Kocak, Jordan G. Bryan, Ranad Humeidi, David Peck, Xiaoyun Wu, Andrew A. Tang, Vickie M. Wang, Sasha A. Bber, Joshua M. Golber, Aravind Subramanian, Eric S. Lander, Gad Getz, Todd R. Golub, and Aviad Tsherniak. Discovering the anti-cancer potential of non-oncology drugs by systematic viability profiling. Nature Cancer, 1:235–248, 2020. doi: 10.1038/s43018-019-0018-6.

Haitham A. Elmarakeby, Justin Hwang, Rand Arafeh, Jett Crowdis, Seetha Gang, David Liu, Saud H. AlDubayan, Keyan Salber, Keegan Korthauer, Stefan Huang, et al. Biologically informed deep neural network for prostate cancer discovery. Nature, 598:348–352, 2021. doi: 10.1038/s41586-021-03922-4.

Octavian-Eugen Ganea, Gary Becigneul, and Thomas Hofmann. Hyperbolic neural networks. In Advances in Neural Information Processing Systems, 2018.

Gene Ontology Consortium. The Gene Ontology knowledgebase in 2023. Genetics, 224(1):iyad031, 2023. doi: 10.1093/genetics/iyad031.

Sonja Hänzelmann, Robert Castelo, and Justin Guinney. GSVA: gene set variation analysis for microarray and RNA-seq data. BMC Bioinformatics, 14:7, 2013. doi: 10.1186/1471-2105-14-7.

Peter M. Haverty, Eva Lin, Jenille Tan, Yihong Yu, Billy Lam, Steve Lianoglou, Audrey Bernal, Daniel P. Strain, Florian Wobbe, David Finkle, et al. Reproducible pharmacogenomic profiling of cancer cell line panels. Nature, 533:333–337, 2016. doi: 10.1038/nature17987.

David Earl Hostallero, Yihui Li, and Amin Emad. Looking beyond the crystal ball: How over-reliance on model accuracy can lead to unreliable drug response prediction in cancer. Nature Communications, 15:1, 2024. doi: 10.1038/s41467-024-52849-5.

Francesco Iorio, Theo A. Knijnenburg, Daniel J. Vis, Graham R. Bignell, Michael P. Menden, Michael Schubert, Nanne Aben, Emanuel Goncalves, Syd Barthorpe, Howard Lightfoot, Thomas Cokelaer, Patricia Greninger, Ewald van Dyk, Han Chang, Heshani de Silva, Holger Heyn, Xianming Deng, Regina K. Egan, Qingsong Liu, Tatiana Mironenko, Xeni Mitropoulos, Laura Richardson, Jinhua Wang, Tinghu Zhang, Sebastian Moran, Sergi Sayols, Maryam Soleimani, David Tamborero, Nuria Lopez-Bigas, Petra Ross-Macdonald, Manel Esteller, Nathanael S. Gray, Daniel A. Haber, Michael R. Stratton, Cyril H. Benes, Lodewyk F. A. Wessels, Julio Saez-Rodriguez, Ultan McDermott, and Mathew J. Garnett. A landscape of pharmacogenomic interactions in cancer. Cell, 166(3):740–754, 2016. doi: 10.1016/j.cell.2016.06.017.

Sarthak Jain and Byron C. Wallace. Attention is not explanation. In Proceedings of the 2019 Conference of the North American Chapter of the Association for Computational Linguistics: Human Language Technologies, pages 3543–3556, 2019. doi: 10.18653/v1/N19-1357.

Jiao Jiang, Xiangyu Sun, Zhenyu Wu, Haoyu Deng, Ziying Luo, Bo Qiang, Feng Zhong, Gen Li, Dongsheng Cao, and Xiangxiang Zeng. DeepTTC: A transformer-based model for predicting cancer drug response. Briefings in Bioinformatics, 23(3):bbac100, 2022. doi: 10.1093/bib/bbac100.

Brent M. Kuenzi, Jisoo Park, Samson H. Fong, Kyle S. Sanchez, John Lee, Jason F. Kreisberg, Jianzhu Ma, and Trey Ideker. DrugCell: A visible neural network to guide precision medicine. Cancer Cell, 38(5):672–684, 2020. doi: 10.1016/j.ccell.2020.09.014.

Qiao Liu, Zhiyong Hu, Rui Jiang, and Meng Zhou. DeepCDR: A hybrid graph convolutional network for predicting cancer drug response. Bioinformatics, 36(Supplement 2):i911–i918, 2020. doi: 10.1093/bioinformatics/btaa822.

Scott M. Lundberg and Su-In Lee. A unified approach to interpreting model predictions. Advances in Neural Information Processing Systems, 30:4765–4774, 2017.

Jianzhu Ma, Michael K. Yu, Samson Fong, Keiichiro Ono, Eric Sage, Barry Demchak, Roded Sharan, and Trey Ideker. Using deep learning to model the hierarchical structure and function of a cell. Nature Methods, 15(4):290–298, 2018. doi: 10.1038/nmeth.4627.

Thanh Nguyen, Thanh-Hoang Nguyen, and Svetha Venkatesh. GraphDRP: A graph neural network for drug response prediction. PLOS ONE, 17(8):e0272820, 2022. doi: 10.1371/journal.pone.0272820.

Maximilian Nickel and Douwe Kiela. Poincare embeddings for learning hierarchical representations. In Advances in Neural Information Processing Systems, 2017.

Michael Schubert, Bertram Klinger, Martina Klünemann, Anja Sieber, Florian Uhlitz, Sascha Sauer, Mathew J. Garnett, Nils Blüthgen, and Julio Saez-Rodriguez. Perturbation-response genes reveal signaling footprints in cancer gene expression. Nature Communications, 9:20, 2018. doi: 10.1038/s41467-017-02391-6.

Thijs S. Stutvoet, Arjan Kol, Elisabeth G. E. de Vries, Mirjam de Bruyn, Rudolf S. N. Fehrmann, Wim Timens, Rinse K. Weersma, Marjolijn N. Lub-de Hooge, Steven de Jong, and Annechien Jorritsma-Smit. MAPK pathway activity plays a key role in PD-L1 expression of lung adenocarcinoma cells. The Journal of Pathology, 249(1):52–64, 2019. doi: 10.1002/path.5280.

Mukund Sundararajan, Ankur Taly, and Qiqi Yan. Axiomatic attribution for deep networks. In International Conference on Machine Learning, pages 3319–3328, 2017.

Sarah Wiegreffe and Yuval Pinter. Attention is not not explanation. In Proceedings of the 2019 Conference on Empirical Methods in Natural Language Processing, pages 11–20, 2019. doi: 10.18653/v1/D19-1002.

Yiheng Zhu, Zhenqiu Ouyang, Wenbo Chen, Ruiwei Feng, Danny Z. Chen, Jian Cao, and Jianhua Wu. TGSA: Protein-protein association-based twin graph neural networks for drug response prediction with similarity augmentation. Bioinformatics, 38(2):461–468, 2022. doi: 10.1093/bioinformatics/btab650.

