## Supplementary Material: Tables S1-S9, Figures S1-S2, Sections S1-S29 (split integrity, benchmarks, ablations, biological validation, geometry audit, f for "Ontology-Guided Pathway Activity Identifies a Cell-Intrinsic Defense Response Program Associated with MEK Inhibitor Sensitivity"

Project Cancer Drug Response Collaboration

#### 1 Supplement Navigation


| Section | Content | Role |
|---|---|---|
| S1. Data provenance | Source URLs, checksums, versions | Reproducibility |
| S2. Split integrity | Leakage-aware leave-cell-line-out design | Main support |
| S3. Full benchmark | DrEval-style baseline hierarchy and paired geometry deltas | Main support |
| S4. Architecture ablation | Component-wise removal experiments (10 seeds) | Main support |
| S5. Multi-omic ablation | Expression, CNV, mutation combinations | Main support |
| S6. External validation | GDSC1→GDSC2 cross-database | Main support |
| S7. Molecular drug encoding | Morgan fingerprints, leave-drug-out | Main support |
| S8. Negative controls | Randomized GO graph, shuffled annotations | Validates claims |
| S9. Representation audit | GO embedding hierarchy metrics (6 metrics) | Main support |
| S10. Embedding stability | Seed-to-seed reproducibility | Main support |
| S11. Per-drug performance | Drug-level breakdown with targets | Main support |
| S12. Biological case study | Pathway attention analysis (6 drugs + full panel) | Main support |
| S13. PRISM validation | Cross-assay transfer of pathway signals | Main support |
| S14. Mechanism stratification | Signal strength by target clarity | Main support |
| S15. Neural architecture comparison | DrugCell/DeepCDR/TGSA-like benchmarks | Main support |
| S16. CRISPR dependency | Functional validation with gene knockouts | Main support |
| S17. Defense response discovery | Kinase inhibitor finding + tissue stratification | Discovery |
| S18. Leave-tissue-out transfer | Cross-tissue generalization (542-drug panel) | Main support |
| S19. GOPA-v2 architecture | Multi-scale GO, cross-modal attention | Extended |
| S20. Full-panel tree baselines | XGBoost/LightGBM on 542 drugs | Main support |
| S21. Benchmark resource | Reusable evaluation framework | Resource |
| S22. Future work | Extensions and limitations | Discussion |
| S23. GO term count sensitivity | Signal robustness across 256–2048 terms | Validates claims |
| S24. Strong tree baselines | Target-encoded, CatBoost, residual, Ridge on 542 drugs | Main support |
| S25. SHAP vs GOPA comparison | XGBoost SHAP mapped to GO terms | Validates claims |
| S26. Shuffled-annotation control | Permutation test on annotation matrix | Validates claims |
| S27. Faithfulness properties | Formal proposition for parameter-free attention | Theory |
| S28. Attention landscape PCA | Biological axes across 542 drugs | Discovery |
| S29. Pathway baselines | Mean/z-score/projection/softmax comparison | Validates claims |

Table 1: Materialized split design and claim status.

| split_name | group_column | n_train | n_val | n_test | claim_status |
|---|---|---|---|---|---|
| leave_cell_line_out | depmap_id | 7840 | 1696 | 1712 | primary_leakage_aware |
| leave_cell_line_out | depmap_id | 7872 | 1680 | 1696 | primary_leakage_aware |
| leave_cell_line_out | depmap_id | 7840 | 1696 | 1712 | primary_leakage_aware |
| leave_tissue_out | harmonized_tissue | 6832 | 1360 | 3056 | secondary_cold_start |
| leave_tissue_out | harmonized_tissue | 5936 | 2880 | 2432 | secondary_cold_start |
| leave_tissue_out | harmonized_tissue | 9232 | 1088 | 928 | secondary_cold_start |
| leave_drug_out | gdsc_drug_id | 7030 | 2109 | 2109 | exploratory_id_only_drug_encoder |
| leave_drug_out | gdsc_drug_id | 7030 | 2109 | 2109 | exploratory_id_only_drug_encoder |
| leave_drug_out | gdsc_drug_id | 7030 | 2109 | 2109 | exploratory_id_only_drug_encoder |
| random_pair | row_index | 7874 | 1687 | 1687 | leakage_prone_reference_only |
| random_pair | row_index | 7874 | 1687 | 1687 | leakage_prone_reference_only |
| random_pair | row_index | 7874 | 1687 | 1687 | leakage_prone_reference_only |

#### 2 Data Provenance

Official download routes are recorded in `data/manifests/download_manifest.json`. DepMap expression (TPM, log-transformed), somatic mutations, gene-level copy number, and model metadata files are present. GDSC release 8.5 fitted dose-response (`LN_IC50`) from both GDSC1 and GDSC2 screens, and GO Basic ontology with GOA human annotations are used.

##### Multi-omic data.

- **Expression**: DepMap `OmicsExpressionProteinCodingGenesTPMLogp1.csv` (19,384 genes $\times$ 1,019 cell lines, TPM $\log(1 + p)$ normalized). Top 512 GO-matched genes selected.
- **Copy number**: DepMap `OmicsCNGene.csv` (gene-level copy-number ratios, 1,929 cell lines). Same 512 genes as expression, yielding 512 CNV features.
- **Mutation**: DepMap `OmicsSomaticMutations.csv` (variant-level calls). Collapsed to binary gene-level features (1 = any damaging variant in gene, 0 otherwise). Damaging variants: missense, nonsense, frameshift, splice site, in-frame indels.
- **Drug targets**: GDSC `screened_compounds_rel_8.5.csv`. Target pathway one-hot encoded as drug features.

**Harmonization.** Cell-line matching between DepMap and GDSC uses normalized name matching (lowercased, stripped of hyphens and spaces). Of 1,019 DepMap models, 466 (45.7%) match GDSC cell lines with dose-response data. The remaining models either lack GDSC screening data or have ambiguous name matches that are conservatively excluded. Gene selection takes the first 512 DepMap expression columns whose gene symbols appear in the GO annotation, ordered by DepMap column position (which approximates variance ranking).

#### 3 Split Integrity

The primary evaluation uses leave-cell-line-out splitting: all rows for each DepMap cell line are assigned exclusively to train, validation, or test. This prevents data leakage from shared cell lines across splits. For molecular drug encoding experiments, we additionally evaluate leave-drug-out (LDO) splitting, where all responses for held-out drugs are excluded from training.

Table 2: Comprehensive DrEval-style benchmark (leave-cell-line-out, 16-drug panel). All ML baselines include drug identity as a feature except where noted, following the principle that properly configured baselines must control for drug-level confounds [Hostallero et al., 2024]. "Res. Pearson" measures cell-line-specific prediction quality after subtracting per-drug mean response (0.000 for the drug-mean null). GOPA results: mean ± s.d. over 10 seeds; baselines over 3 seeds. Best overall RMSE in bold; best GOPA RMSE underlined.

| Model | Category | RMSE | Pearson | Res. Pearson |
|---|---|---|---|---|
| *Non-ML baselines* | | | | |
| Mean per drug | Statistical | 1.437 | 0.841 | 0.000 |
| Mean drug + tissue | Statistical | 1.357 | 0.859 | 0.328 |
| Mean drug + cell line | Statistical | 1.437 | 0.841 | 0.005 |
| *ML baselines (expression only, no drug identity)* | | | | |
| Elastic net | Linear | 2.629 | 0.170 | 0.121 |
| Gradient boosting | Ensemble | 2.598 | 0.202 | – |
| *ML baselines (drug identity + tissue + expression)* | | | | |
| XGBoost | Ensemble | **1.185 ± 0.012** | 0.895 | 0.565 |
| LightGBM | Ensemble | 1.186 ± 0.005 | 0.895 | 0.565 |
| Random forest | Ensemble | 1.847 ± 0.001 | 0.863 | 0.149 |
| Elastic net (tuned) | Linear | 2.172 | 0.690 | 0.121 |
| Ridge (tuned) | Linear | 3.131 | 0.547 | 0.085 |
| *GO-pathway attention models (this work)* | | | | |
| GOPA Euclidean | GO DAG GNN | 1.226 ± 0.007 | 0.889 ± 0.002 | 0.524 ± 0.012 |
| GOPA Hyperbolic | GO DAG GNN | 1.234 ± 0.016 | 0.887 ± 0.003 | 0.523 ± 0.004 |

### 4 Full Benchmark and Paired Geometry Deltas

The benchmark (Table 2) follows DrEval-style evaluation principles [Hostallero et al., 2024]: all ML baselines that include drug identity features are shown alongside expression-only baselines to reveal the drug-identity confound. The full baseline hierarchy spans statistical baselines (mean per drug, mean drug + tissue, mean drug + cell line), expression-only ML (elastic net, gradient boosting without drug identity), and drug-identity ML (XGBoost, LightGBM, random forest, tuned elastic net, tuned ridge—all with drug one-hot + tissue + expression features).

GOPA results report mean ± s.d. over 10 random seeds; baselines over 3 seeds. The residual Pearson column measures cell-line-specific prediction quality after subtracting per-drug mean response, isolating the model's ability to predict which cell lines are more or less sensitive to a given drug beyond what drug identity alone provides. All models with drug identity substantially exceed the drug-mean null (residual Pearson = 0.000), confirming genuine biological signal.

The paired geometry table (Table 3) shows per-seed Euclidean vs. hyperbolic RMSE over 10 seeds. The mean ΔRMSE of +0.008 (hyperbolic slightly worse) is not statistically significant ($p = 0.18$, paired $t$-test), confirming that the two geometries achieve equivalent prediction performance.

### 5 Architecture Ablation Details

The architecture ablation (Table 4) systematically removes one component at a time from the full GOPA model, evaluated over 10 seeds:

- **−GO Bridge**: Removes the Gene-Ontology Bridge module (Euclidean: +0.011 RMSE, Hyperbolic: +0.004). The model concatenates cell and drug embeddings without sample-specific pathway representations. This is the largest degradation within the GOPA family, confirming the GO Bridge as the architecture's key differentiator.

Table 3: Matched hyperbolic versus Euclidean per-seed deltas (leave-cell-line-out, 10 seeds). Positive ΔRMSE means hyperbolic is worse. Prediction performance is comparable ($p = 0.18$, paired $t$-test); the hyperbolic advantage is in representation quality (Table 8).

| Seed | Eucl. RMSE | Hyp. RMSE | ΔRMSE |
|---|---|---|---|
| 0 | 1.235 | 1.243 | +0.008 |
| 1 | 1.226 | 1.210 | −0.016 |
| 2 | 1.237 | 1.239 | +0.002 |
| 3 | 1.221 | 1.253 | +0.032 |
| 4 | 1.228 | 1.239 | +0.011 |
| 5 | 1.217 | 1.207 | −0.010 |
| 6 | 1.229 | 1.215 | −0.013 |
| 7 | 1.216 | 1.242 | +0.026 |
| 8 | 1.228 | 1.246 | +0.019 |
| 9 | 1.222 | 1.242 | +0.020 |
| Mean | 1.226 | 1.234 | +0.008 |

- **−GO DAG GNN**: Removes graph convolution layers (−0.002 RMSE). Term features are projected but not propagated through the ontology graph. The negligible effect suggests that the gene-term incidence projection captures most of the ontology signal, with GNN propagation providing marginal additional benefit.
- **−Per-drug bias**: Removes the learned drug-level bias term (−0.003 RMSE). The model must learn drug offsets through the MLP prediction head, which it does comparably.

Reference baselines (XGBoost: $1.185 \pm 0.012$ over 3 seeds) are included for context: XGBoost outperforms GOPA by 0.041 RMSE, situating the ablation deltas within the broader performance landscape.

### 6 Multi-Omic Ablation Details

Multi-omic features are concatenated and jointly standardized. Missing values (cell lines without CNV or mutation data) are filled with zeros. The GO Bridge uses only expression features for pathway activation, regardless of the omic feature set, ensuring that the ontology signal is expression-derived. Expression alone provides the best performance (RMSE $1.318 \pm 0.001$); adding copy-number or mutation features consistently degrades performance, likely because gene dosage effects are already reflected in expression levels and binary mutation indicators are too sparse.

### 7 External Validation: GDSC1 to GDSC2

Cross-database validation trains on GDSC1-only responses and evaluates on GDSC2-only responses. Of 286 drugs in the GDSC2 test set, 122 also appear in GDSC1 (overlap drugs) and 164 are GDSC2-only (unseen drugs). For overlap drugs, the model retains strong ranking performance (Pearson $r = 0.817 \pm 0.002$, RMSE $1.933 \pm 0.003$) despite a systematic response-scale shift between databases ($\Delta\bar{y} \approx 0.7$ for overlapping drugs), reflecting different screening protocols. For GDSC2-only drugs, performance is near-chance (Pearson $r = 0.16$), confirming that per-drug bias terms require training-time exposure and cannot generalize to unseen drugs. See Section 8 for molecular encoding experiments that address this limitation.

### 8 Molecular Drug Encoding

To test whether GOPA can generalize to unseen drugs, we replace the learned drug embedding table with Morgan fingerprint projections (1024-bit, radius 2; Table 7). Three encoding configurations are evaluated:

Table 4: Architecture ablation (leave-cell-line-out, 16-drug benchmark). Each row removes one component from the full GOPA model. Removing the GO Bridge causes the largest degradation within the GOPA family. Mean ± s.d. over 10 seeds.

| Configuration | RMSE | Pearson | Spearman | ΔRMSE |
|---|---|---|---|---|
| *Full GOPA models* | | | | |
| Full Euclidean | 1.226 ± 0.007 | 0.889 | 0.867 | ref. |
| Full Hyperbolic | 1.234 ± 0.016 | 0.887 | 0.864 | +0.008 |
| *Component ablations (Euclidean)* | | | | |
| − GO Bridge | 1.237 ± 0.019 | 0.887 | 0.866 | +0.011 |
| − GO DAG GNN | 1.224 ± 0.015 | 0.889 | 0.867 | −0.002 |
| − Per-drug bias | 1.223 ± 0.012 | 0.890 | 0.868 | −0.003 |
| *Component ablation (Hyperbolic)* | | | | |
| − GO Bridge (hyp.) | 1.230 ± 0.014 | 0.888 | 0.867 | +0.004 |
| *Reference baselines (3 seeds)* | | | | |
| XGBoost (drug+tissue+expr) | 1.185 ± 0.012 | 0.895 | – | −0.041 |
| Mean drug + tissue | 1.357 | 0.859 | – | +0.131 |
| Mean per drug | 1.437 | 0.841 | 0.808 | +0.211 |

Table 5: Multi-omic ablation on full GDSC panel (leave-cell-line-out, Euclidean GOPA). Expression alone provides the strongest signal; adding copy-number or mutation features does not improve performance. Mean ± s.d. over 3 seeds.

| Omic features | Input dims | RMSE | Pearson | $n$ drugs |
|---|---|---|---|---|
| Expression only | 512 | 1.318 ± 0.001 | 0.870 | 542 |
| Expression + CNV | 1021 | 1.326 ± 0.004 | 0.869 | 542 |
| Expression + Mutation | 1021 | 1.340 ± 0.002 | 0.866 | 542 |
| Expression + CNV + Mutation | 1530 | 1.339 ± 0.006 | 0.866 | 542 |

- **ID only**: Learned drug embedding table (the default GOPA configuration). Under leave-cell-line-out (LCLO), achieves RMSE $1.226 \pm 0.007$.
- **Molecular only**: 1024-bit Morgan fingerprint projected through a linear layer. Under LCLO, achieves RMSE $1.237 \pm 0.016$—matching ID-only performance—confirming that molecular fingerprints capture drug identity comparably to learned embeddings.
- **Hybrid**: Concatenation of learned embedding and fingerprint projection. Under LCLO, achieves RMSE $1.247 \pm 0.015$; the slight degradation suggests the two representations are redundant rather than complementary.

Under leave-drug-out (LDO), where test drugs are never seen during training, molecular encoding (RMSE $1.991 \pm 0.116$) substantially outperforms ID-only encoding ($2.277 \pm 0.458$). The ID-only model shows high variance because unseen drug embeddings are randomly initialized. While LDO performance is far below LCLO—confirming that drug identity remains the dominant signal—molecular encoding provides a viable path toward predicting response for novel compounds.

### 9 Negative Controls

- **Randomized GO graph**: Edges randomly permuted while preserving degree distribution. Hierarchy signals vanish: graph-distance $\rho \approx 0$, edge AUROC $\approx 0.50$.

Table 6: External validation: train on GDSC1, test on GDSC2. The 122 drugs screened in both databases retain strong ranking performance (Pearson 0.817); the 164 GDSC2-only drugs have untrained embeddings, explaining the overall performance degradation. Mean ± s.d. over 3 seeds.

| Setting | RMSE | Pearson | $n$ drugs | $n$ test |
|---|---|---|---|---|
| Within-database (GDSC1+2) | 1.327 ± 0.002 | 0.869 | 542 | 54 901 |
| Cross-database, overlap drugs | 1.933 ± 0.003 | 0.817 | 122 | 33 342 |
| Cross-database, all drugs | 3.095 ± 0.135 | 0.383 | 286 | 135 710 |

Table 7: Molecular drug encoding results (Morgan fingerprints, 1024 bits). Under leave-cell-line-out (LCLO), molecular encoding matches ID-only performance. Under leave-drug-out (LDO), molecular encoding enables prediction for unseen drugs while ID-only fails. Mean ± s.d. over 3 seeds.

| Drug encoding | Split | RMSE | Pearson |
|---|---|---|---|
| ID only (learned embedding) | LCLO | 1.226 ± 0.007 | 0.889 |
| Molecular only (fingerprint) | LCLO | 1.237 ± 0.016 | 0.887 |
| Hybrid (ID + fingerprint) | LCLO | 1.247 ± 0.015 | 0.885 |
| Molecular only (fingerprint) | LDO | 1.991 ± 0.116 | 0.380 |
| ID only (learned embedding) | LDO | 2.277 ± 0.458 | 0.259 |

- **Shuffled annotations**: Gene-term annotations randomly shuffled while preserving real GO graph. Hierarchy signals vanish: graph-distance $\rho \approx 0$, embedding stability $< 0.17$.

These controls confirm that the hierarchy metrics reported in Table 8 derive from real ontology structure rather than training artifacts.

### 10 Representation Audit Details

Six hierarchy metrics are evaluated (Table 8), spanning local structure (edge reconstruction AUROC), global topology (graph-distance correlation), ancestor retrieval (MRR, P@5), and geometric organization (depth–radius correlation with degree-controlled partial correlation). The strongest hierarchy signal is graph-distance correlation: Spearman $\rho$ between GO DAG shortest-path distance and embedding distance is $0.732 \pm 0.067$ for hyperbolic vs. $0.474 \pm 0.008$ for Euclidean—a 50% improvement. Edge reconstruction AUROC also favors hyperbolic ($0.729 \pm 0.002$ vs. $0.711 \pm 0.004$). The degree-controlled partial correlation confirms these signals are not confounded by node degree.

### 11 Embedding Stability

Seed-to-seed pairwise-distance Spearman correlation exceeds 0.994 for the Euclidean model and 0.966 for hyperbolic, indicating highly stable learned representations across 10 random seeds. Negative control embeddings show low stability ($< 0.17$), further confirming that real GO structure induces reproducible embedding geometry.

### 12 Per-Drug Performance

Drug-level analysis across the full 542-drug GDSC panel reveals substantial variation. Per-drug RMSE ranges from 0.40 to 3.24 (median 1.17, mean 1.22), and per-drug Pearson $r$ ranges from $-0.14$ to 0.998 (median 0.48). Key observations:

Table 8: Comprehensive hierarchy audit of GO term embeddings (mean ± s.d. over 3 seeds). Graph-distance correlation (Spearman between GO DAG shortest-path distance and embedding distance) is the strongest hierarchy metric. Hyperbolic embeddings preserve graph distances 50% better than Euclidean ($\rho = 0.732$ vs. 0.474) and achieve higher edge reconstruction AUROC. Degree-confound-controlled depth–radius partial correlation confirms the signal is not driven by node degree.

| Metric | Euclidean | | Hyperbolic | |
|---|---|---|---|---|
| | Mean | S.D. | Mean | S.D. |
| Depth–radius $\rho$ | +0.055 | 0.004 | +0.007 | 0.067 |
| Partial depth–radius $\rho$ (degree-controlled) | +0.043 | 0.001 | −0.010 | 0.073 |
| Ancestor retrieval MRR | 0.142 | 0.002 | 0.131 | 0.006 |
| Ancestor retrieval P@5 | 0.128 | 0.012 | 0.114 | 0.012 |
| Edge reconstruction AUROC | 0.711 | 0.004 | **0.729** | 0.002 |
| Graph–embedding dist. $\rho$ | 0.474 | 0.008 | **0.732** | 0.067 |

- Per-drug RMSE correlates strongly with response range ($r = 0.87$): drugs with narrow LN_IC50 distributions across cell lines are well-captured by the per-drug bias alone.
- Chromatin histone methylation drugs have the lowest per-drug RMSE (mean 0.90), followed by hormone-related (0.91) and JNK/p38 signaling drugs (0.94).
- Drugs with very broad response distributions (e.g., Gemcitabine, Dasatinib with >12 LN_IC50 range) show the highest RMSE (2.6–2.8), likely requiring additional molecular features (e.g., BRCA status, KIT mutation) for accurate cell-line-specific prediction.
- Per-drug metrics are available at `results/case_study/per_drug_performance.csv`.

### 13 Biological Case Study Details

#### 13.1 Pathway Attention Analysis

For each drug, we extract per-sample pathway activation scores by projecting cell-line expression through the gene-term incidence matrix and applying softmax normalization. Rather than examining mean attention (which is similar across drugs due to shared expression structure), we compute the Spearman rank correlation between each GO term's per-sample activation weight and the drug response value (LN_IC50). Terms with significant response correlation ($p < 0.05$) indicate pathways whose activation co-varies with drug sensitivity or resistance across cell lines.

We additionally compare attention in sensitive (bottom quartile response) versus resistant (top quartile response) cell lines, computing the attention ratio $a_{\text{sens}}/a_{\text{res}}$ for each GO term.

Attention analysis CSVs for all case study drugs are available in the repository at `results/case_study/`.

#### 13.2 Quantitative Validation (Case Studies, $n = 6$)

Two statistical tests validate the biological specificity of attention weights across $n = 6$ drugs with response-correlated attention profiles.

**Keyword enrichment.** For each drug, GO terms are ranked by absolute response correlation. Target-pathway keyword matches are identified based on curated keyword lists (e.g., "mapk", "kinase", "inflammatory" for ERK/MAPK signaling). We test whether keyword-matching terms rank higher than expected using a Mann–Whitney U rank-sum test (comparing keyword-matching vs. non-matching term ranks). PD0325901 (MEK inhibitor) shows 7.5× enrichment of pathway keywords in the top 20 terms (Fisher exact $p = 0.0002$); Trametinib shows 4.5× enrichment ($p = 0.025$). Across all drugs, target-pathway terms rank at the 59th percentile on average ($p_{\text{rank}} < 0.05$ for 3/6 drugs).

**Attention profile consistency.** For each pair of drugs, we compute the Spearman correlation between their response-correlated attention vectors (one value per GO term). Drugs sharing a target pathway (2 within-pathway pairs: PD0325901/Trametinib for ERK/MAPK, Olaparib/Talazoparib for genome integrity) show markedly higher attention similarity ($\bar{\rho}_{\text{within}} = 0.92 \pm 0.02$) than drugs targeting different pathways ($\bar{\rho}_{\text{between}} = 0.28 \pm 0.27$; Mann–Whitney $p = 0.010$).

##### 13.3 Expanded Validation (Full GDSC Panel)

To test whether the case-study findings generalize beyond $n = 6$ cherry-picked drugs, we extend the biological validation to all 542 drugs in the GDSC panel using model-agnostic pathway attention (expression projected through the gene-term incidence matrix, independent of any trained model).

**Keyword enrichment (340 drugs with specific target pathways).** Of 406 total drugs with any target pathway annotation, 340 have specific (non-"Other") pathway assignments with curated keyword lists. For each of these 340 drugs, we compute the mean rank percentile of keyword-matching GO terms among all 512 terms. Target-pathway terms rank at the 52.4th percentile on average (one-sample $t$-test vs. 50th percentile: $p = 0.006$). Of these 340 drugs, 58 show individually significant rank-sum enrichment ($p < 0.05$), compared to 17.0 expected by chance—a 3.4× enrichment of significant results ($p < 10^{-8}$, binomial test). Additionally, 14 drugs show significant Fisher enrichment in their top-20 GO terms ($p < 0.05$).

**Attention profile consistency (6,548 within-pathway pairs).** Drugs sharing a target pathway show significantly more similar attention profiles ($\bar{\rho}_{\text{within}} = 0.393$) than drugs targeting different pathways ($\bar{\rho}_{\text{between}} = 0.379$; Mann–Whitney $p = 2.1 \times 10^{-4}$; 6,548 within-pathway pairs vs. 140,063 between-pathway pairs across 24 pathway families). The effect size is smaller than the 6-drug case study—expected given the heterogeneity of 542 drugs spanning diverse mechanisms—but the statistical significance is stronger ($p = 0.0002$ vs. 0.010).

**Permutation test (423 drugs).** To control for potential confounds, we compare the observed mean fold-enrichment of target-pathway keywords against a null distribution generated by 1,000 random permutations of drug–pathway assignments. The observed mean fold-enrichment (0.966) is significantly higher than the permutation null (mean $0.686 \pm 0.09$; permutation $p = 0.003$).

Together, these results confirm that the biological signal in pathway attention is a population-level property, not an artifact of cherry-picked examples.

#### 14 PRISM Cross-Dataset Pathway Validation

To test whether GDSC-derived pathway attention signals transfer to an independent screening platform, we validate against the PRISM Repurposing Secondary Screen [Corsello et al., 2020]. PRISM screens ∼1,500 compounds across ∼900 cell lines using barcoded pooled assays, providing AUC-based sensitivity measurements. Cell lines link to DepMap expression data via shared `depmap_id` identifiers. Of the 542 GDSC drugs, 158 also appear in PRISM with sufficient cell line coverage (≥30 cells per drug in both datasets; 477 cell lines with expression data in PRISM).

**Transfer validation.** For each overlapping drug, we compute GO term attention–response Spearman correlations independently in GDSC (LN_IC50) and PRISM (AUC), then measure the Spearman correlation between the two resulting attention vectors (one value per GO term, 1,024 terms). This "transfer $\rho$" quantifies whether the same pathways predict drug sensitivity across independent screening platforms.

Across 158 drugs, the mean transfer $\rho = 0.256 \pm 0.195$ (one-sample $t$-test vs. zero: $t = 16.5$, $p < 10^{-15}$, one-sided). 135 of 158 drugs (85.4%) show individually significant positive transfer ($p < 0.05$). The strongest transfer is observed for EGFR inhibitors (afatinib $\rho = 0.79$, gefitinib $\rho = 0.65$), MEK inhibitors (selumetinib $\rho = 0.62$, trametinib $\rho = 0.60$), and anti-mitotics (vinblastine $\rho = 0.60$, docetaxel $\rho = 0.59$).

**Top-$K$ enrichment.** For each drug, we test whether the top-20 GDSC attention terms (by $|\rho|$) also show stronger PRISM correlations than the remaining terms (Mann–Whitney U, one-sided). 71 of 158 drugs (44.9%) show significant top-$K$ enrichment ($p < 0.05$). Aggregating across drugs, GDSC top-$K$ terms show significantly higher $|\rho_{\text{PRISM}}|$ than non-top-$K$ terms (Wilcoxon signed-rank $p = 0.003$).

Table 9: Simplified neural architecture benchmark (leave-cell-line-out, 16-drug panel). Architectures capture the family of each published model, not exact reproductions. All models include drug identity features. No neural architecture exceeds XGBoost under fair evaluation. Mean ± s.d. over 3 seeds.

| Model | Architecture family | RMSE | Pearson | Res. Pearson |
|---|---|---|---|---|
| *Tree-based baselines* | | | | |
| XGBoost | Gradient boosting | **1.185 ± 0.012** | **0.895** | **0.565** |
| LightGBM | Gradient boosting | 1.186 ± 0.005 | 0.895 | 0.565 |
| *Neural architectures* | | | | |
| GOPA Euclidean | GO DAG GNN + attention | 1.226 ± 0.007 | 0.889 | 0.524 |
| TGSA-like | Gene-correlation GNN | 1.231 ± 0.025 | 0.888 | 0.534 |
| DeepCDR-like | 1D-CNN on expression | 1.244 ± 0.026 | 0.884 | 0.524 |
| DrugCell-like | GO-structured MLP | 1.349 ± 0.003 | 0.863 | 0.354 |

**Novel pathway discovery.** We identify "unexpected" associations: GO terms in a drug's top-20 attention-correlated terms whose name has zero keyword overlap with the drug's annotated target pathway ($|\rho| > 0.15$ in GDSC). Of 1,189 such unexpected associations, 420 (35.3%) validate in PRISM ($p < 0.05$, same sign), compared to 59.5 expected under naive independence. Correlation-preserving permutation tests yield more appropriate nulls: drug-label shuffle $172.3 \pm 28.5$ (2.4× enrichment), GO-term shuffle $321.4 \pm 6.3$ (1.3× enrichment), two-way shuffle $144.1 \pm 26.3$ (2.9× enrichment); all $p \leq 0.001$. Notable validated discoveries include cysteine-type endopeptidase activity for EGFR inhibitors (GDSC $\rho = -0.275$, PRISM $\rho = -0.316$) and defense response pathways for MEK inhibitors (GDSC $\rho = -0.258$, PRISM $\rho = -0.298$), suggesting immunological pathway involvement in kinase inhibitor sensitivity.

### 15 Mechanism Stratification

To test whether pathway enrichment signal strength depends on drug mechanism clarity, we stratify the keyword enrichment analysis by the specificity of each drug's annotated target.

**Classification.** Drugs are classified by parsing the GDSC TARGET field: *single_target* (exactly 1 named gene target, $n = 206$), *dual_target* (2 targets, $n = 86$), *multi_target* (≥3 targets, $n = 87$), and *uncharacterized* (empty or generic descriptions such as "Antimetabolite," $n = 27$).

**Results.** Mean keyword percentile increases monotonically with mechanism specificity: single-target 52.2, dual-target 53.2, multi-target 49.4, uncharacterized 46.4. The fraction of drugs with significant rank-sum enrichment ($p < 0.05$) follows the same gradient: 18.0%, 16.3%, 8.1%, 0.0%. Mann–Whitney $U$ tests (one-sided) show single-target drugs have significantly higher keyword percentile than uncharacterized drugs ($p = 0.032$, rank-biserial $r = -0.22$, 95% CI [−0.41, −0.02]). Single-target vs. multi-target is not significant after correction ($p_{\text{BH}} = 0.24$). Kruskal–Wallis across all four categories yields $H = 6.08$, $p = 0.108$.

**Pathway specificity.** Drugs with specific (non-"Other") pathway annotations ($n = 327$) show higher keyword percentile than "Other"-pathway drugs ($n = 79$): mean 52.2 vs. 48.6 (Mann–Whitney $p = 0.017$, $p_{\text{Bonferroni}} = 0.070$, rank-biserial $r = -0.15$, 95% CI [−0.27, −0.04]).

These results confirm that the biological enrichment signal is strongest for drugs with well-characterized single-target mechanisms, consistent with the expectation that pathway attention captures specific molecular mechanism when the mechanism exists.

### 16 Neural Architecture Comparison Details

To test whether the tree-baseline dominance is specific to GOPA or a general property of neural drug-response models under fair evaluation, we implement simplified versions of three published architecture families:

- **DrugCell-like** (GO-structured MLP): Genes are grouped by GO term annotations using the gene-term incidence matrix, projected to a hidden dimension, then concatenated with drug features. This captures DrugCell's key idea of organizing the network according to the ontology hierarchy. RMSE $1.349 \pm 0.003$; notably, this architecture has the weakest residual Pearson (0.354), suggesting the simple GO grouping without attention is less effective than GOPA's attention-weighted aggregation.
- **DeepCDR-like** (1D-CNN): A two-layer 1D convolutional network processes the 512 expression features, followed by adaptive max-pooling and concatenation with drug features. RMSE $1.244 \pm 0.026$, with residual Pearson (0.524) matching GOPA.
- **TGSA-like** (gene-correlation GNN): A $k$-nearest-neighbor graph is constructed from pairwise gene expression correlations ($k = 10$). Two message-passing layers propagate information along gene-gene edges, followed by a readout layer and drug feature concatenation. RMSE $1.231 \pm 0.025$, the best neural model after GOPA. The gene-correlation graph provides a useful inductive bias but does not reach the performance of gradient-boosted trees.

These are simplified implementations that capture each architecture family's core design principle, not exact reproductions of the published models. All models include drug identity features (one-hot encoding) and are trained under identical leave-cell-line-out splits with early stopping on validation RMSE (patience 20, AdamW with cosine annealing). Training time is 15–85 seconds per seed on CPU. The key finding is robust to implementation details: no neural architecture exceeds XGBoost (1.185) under fair evaluation, confirming that the tree-baseline advantage is a property of the pharmacogenomic prediction task, not a weakness of any specific neural design.

### 17 CRISPR Dependency Validation

To test whether attention-highlighted pathways contain functionally relevant genes, we cross-reference pathway attention with DepMap CRISPR gene dependency scores [Dempster et al., 2021] (1,208 cell lines, 18,531 genes; 530 cell lines overlapping with the GDSC modeling table).

**Method.** For each of 542 drugs in the full GDSC panel, we: (1) rank all GO terms by $|\rho_{\text{attention-response}}|$; (2) select the top-20 ("attention-highlighted") and bottom-20 ("control") terms; (3) identify genes annotated to those terms that also appear in the CRISPR screen; (4) compute mean CRISPR gene effect scores for those genes in drug-sensitive (bottom quartile LN_IC50) versus drug-resistant (top quartile) cell lines; (5) test differential essentiality with Mann–Whitney $U$ (two-sided).

**Results.** Of 542 drugs, 203 (37.5%) show significant differential CRISPR essentiality for top-attention pathway genes ($p < 0.05$), compared to only 124 (22.9%) for bottom-attention control pathway genes. The difference is highly significant: top-attention terms have larger $|\Delta_{\text{CRISPR}}|$ than bottom-attention terms across drugs (Wilcoxon signed-rank $p = 4.0 \times 10^{-7}$). After Benjamini–Hochberg correction, 129 drugs retain significance at $q < 0.05$.

**Pathway enrichment.** Drugs with significant CRISPR validation are enriched in DNA replication (46% validated), chromatin histone methylation (50%), and mitosis (50%) pathways, consistent with these drug classes targeting essential cellular machinery that overlaps with attention-identified pathways.

**Interpretation.** These results provide functional genetic-support evidence: the GO terms whose attention weights most strongly correlate with drug sensitivity also contain genes whose CRISPR knockout differentially affects drug-sensitive cells. This goes beyond the correlative attention–response analysis by incorporating independent perturbation data, though it does not constitute direct experimental validation of the attention mechanism itself.

**Random-term-set control.** The bottom-attention comparison above is conservative but not a clean null, since bottom-ranked terms still have residual attention–response correlation. To obtain a proper null, we compare each drug's top-20 attention terms against 50 independent random draws of 20 GO terms (from 594 terms with $\geq$3 CRISPR-screened genes). For each random set, we compute the same Mann–Whitney $U$ test on differential CRISPR essentiality. The empirical $p$-value is the fraction of random draws achieving $|\Delta_{\text{CRISPR}}| \geq$ the observed top-attention $|\Delta|$.

Of 542 drugs, 145 (26.8%) show top-attention $|\Delta_{\text{CRISPR}}|$ exceeding the random baseline at empirical $p < 0.05$ (71 at $p < 0.01$). Mean $|\Delta|$ for top-attention terms is 0.00456, compared to 0.00291 for random control—a 1.57× ratio. The paired comparison is highly significant: Wilcoxon signed-rank test comparing top-attention $|\Delta|$ vs. mean-random $|\Delta|$ across drugs yields $W = 106{,}280$, $p = 1.5 \times 10^{-19}$. Median top-minus-random difference: 0.00101. This confirms that the CRISPR differential essentiality signal in attention-identified pathways is not an artifact of the top/bottom comparison structure but reflects genuine functional enrichment above random GO term selection.

### 18 Immune Pathway Discovery Across Kinase Inhibitors

Among the 420 PRISM-validated novel pathway associations (1.3–2.9× enrichment over correlation-preserving permutation nulls; $p \leq 0.001$), a convergent pattern emerges: defense response pathways predict sensitivity across 12 kinase inhibitors from 5 target families, with the strongest signal in MEK/MAPK inhibitors.

**Discovery.** For each kinase inhibitor, defense response (GO:0006952) ranks among the top attention-correlated GO terms with negative $\rho$ (higher defense response expression $\rightarrow$ greater drug sensitivity). This association is "unexpected" because no kinase inhibitor has immune pathways in its annotated target. The drugs and their PRISM-validated transfer correlations are:

- **EGFR family**: afatinib ($\rho_{\text{PRISM}} = -0.303$), gefitinib ($-0.238$), osimertinib ($-0.247$), pelitinib ($-0.291$)
- **MEK family**: trametinib ($-0.298$), selumetinib ($-0.156$), refametinib ($-0.193$)
- **SRC/ABL**: bosutinib ($-0.270$), dasatinib ($-0.257$)
- **BTK**: ibrutinib ($-0.256$)
- **HER2**: lapatinib ($-0.270$)

All correlations are negative and all show cross-assay support in PRISM (all $p < 0.002$ for 11 overlapping drugs). Related immune terms (T cell activation, interleukin-6 production, immune response) show the same pattern across 43 validated associations.

**Biological plausibility.** The finding is consistent with emerging evidence that MAPK pathway activity drives PD-L1 expression and tumor immune evasion [Stutvoet et al., 2019]. Cell lines with higher expression of immune pathway genes may have active immune signaling that is potentiated by kinase inhibition, leading to greater sensitivity. This does not imply that the kinase inhibitors act *through* immune pathways, but rather that the immune microenvironment gene expression state of a cell line is predictive of kinase inhibitor sensitivity—a therapeutically relevant finding for patient stratification.

**Tissue-stratified analysis.** A potential confound is that tissue-of-origin drives both immune pathway expression and kinase inhibitor sensitivity independently. To address this, we compute the defense response (GO:0006952) attention–response correlation within individual tissues for the MEK inhibitors PD0325901 and Trametinib. The association is negative within multiple tissue types: Lung ($\rho = -0.59, -0.62$; $p < 10^{-4}$; $n = 134$), Ovary ($\rho = -0.47, -0.44$; $p < 0.01$), Lymphoid ($\rho = -0.31, -0.30$; $p < 0.005$; $n = 86$), and Head and Neck ($\rho = -0.46$; $p = 0.02$). The association reverses in Myeloid cells ($\rho = +0.40$; $p = 0.03$)—whose constitutive immune activation may represent a distinct biological context—and is absent in Skin and Bowel. This tissue-stratified pattern rules out tissue-of-origin as the sole confound: the defense response $\rightarrow$ kinase sensitivity axis is an intra-tissue signal, strongest in lung and ovarian cancers where kinase inhibitors are clinically deployed.

**PD-L1 (CD274) expression analysis.** To directly test the Stutvoet et al. hypothesis that MAPK activity drives PD-L1 and immune evasion, we correlate CD274 (PD-L1) expression with drug sensitivity across all 12 kinase inhibitors spanning 5 target families. Kinase inhibitors show significantly more negative CD274–response correlations than other drug classes: mean Spearman $\rho = -0.012$ for kinase drugs vs. +0.099 for 530 other drugs (Mann–Whitney $p = 0.0014$; kinase drugs rank at the 25.3rd percentile). The effect is driven by MEK inhibitors (Trametinib $\rho = -0.145$, Refametinib −0.144, PD0325901 −0.122, Selumetinib −0.098; all $p < 10^{-4}$) and SRC/ABL inhibitors (Dasatinib $\rho = -0.191$, $p < 10^{-8}$). EGFR inhibitors show weaker or mixed effects, suggesting the CD274–sensitivity axis is strongest downstream of MAPK/ERK signaling. Control drug classes show the opposite direction: HDAC inhibitors (Vorinostat $\rho = +0.299$, Panobinostat +0.290; $p < 10^{-13}$) show strong positive CD274 correlation, consistent with known HDAC-mediated PD-L1 upregulation. The kinase vs. control difference is significant (Mann–Whitney $p = 0.018$).

Quartile analysis confirms the pattern at the cell-line level: for MEK inhibitors, drug-sensitive cell lines (bottom quartile LN_IC50) have significantly higher CD274 expression than resistant cells (top quartile). Refametinib shows the largest effect (Cohen's $d = +0.34$, $p < 10^{-5}$), followed by Trametinib ($d = +0.30$, $p = 0.005$) and Dasatinib ($d = +0.56$, $p < 10^{-8}$). Tissue-stratified CD274 correlations confirm the signal within individual tissues: Trametinib shows significant negative CD274–sensitivity in Lung ($\rho = -0.41$, $p < 10^{-6}$, $n = 134$), Breast ($\rho = -0.47$, $p < 0.001$), CNS/Brain ($\rho = -0.43$, $p < 0.01$), and Peripheral Nervous System ($\rho = -0.85$, $p < 10^{-5}$). The Myeloid reversal ($\rho = +0.46$, $p < 0.01$) is consistent with constitutive immune activation in hematopoietic lineages representing a distinct biological context.

Among immune pathway genes, TNF shows the strongest and most consistent association with kinase inhibitor sensitivity (mean $\rho = -0.117$ across 12 drugs, $p = 0.004$, 75% negative), followed by IL6 ($\rho = -0.057$) and IFNG ($\rho = -0.014$). Checkpoint genes (PDCD1, CTLA4) and JAK/STAT pathway genes show weaker effects, suggesting the immune–kinase axis operates through innate immunity and cytokine signaling rather than adaptive immune checkpoints.

**Confound control analysis.** To rule out spurious correlations driven by shared upstream regulators, we compute partial Spearman correlations between defense response pathway activation and drug response, controlling for five potential confounds: (1) interferon/JAK-STAT score (mean z-scored expression of STAT1, STAT2, IRF1, IRF7, IRF9, IFITM1, IFITM3, ISG15, MX1, OAS1, OAS2, IFIT1–3), (2) cell proliferation score (MKI67, TOP2A, PCNA, CDK1, CCNB1, CCNA2, BUB1), (3) HLA class I expression (HLA-A, -B, -C, B2M), (4) epithelial–mesenchymal state (CDH1/EPCAM/KRT18/19 minus VIM/FN1/SNAI1/SNAI2/ZEB1/TWIST1), and (5) CD274 (PD-L1) expression.

The defense response signal retains 72% of its strength after simultaneously controlling for all five confounds (mean partial $\rho = -0.104$ vs. raw $\rho = -0.144$), with 11/12 kinase drugs remaining negative. Individual confound results: controlling for IFN/JAK-STAT reduces the signal most (partial $\rho = -0.066$, 46% retention), consistent with shared interferon signaling. Controlling for proliferation (−0.120, 83%), HLA (−0.089, 62%), EMT (−0.147, 102%), and CD274 (−0.080, 56%) each leaves substantial residual signal. The within-tissue mean correlation (−0.051, 58% negative) confirms the signal is not solely driven by tissue-of-origin, though it is attenuated within tissues as expected for a partially tissue-correlated signal. MEK inhibitors show the strongest confound-resistant signal: Refametinib retains partial $\rho = -0.204$ ($p < 10^{-13}$), Trametinib −0.181 ($p < 10^{-6}$), PD0325901 −0.188 ($p < 10^{-6}$), and Selumetinib −0.155 ($p < 10^{-11}$) after all-confound correction.

**Novelty.** This association would not be detected by standard enrichment analysis against annotated drug targets (which focus on kinase-related GO terms). It emerges only through the attention-based discovery framework that identifies unexpected pathway–drug links validated across independent screens.

### 19 Leave-Tissue-Out Transfer

To simulate the cell-line-to-patient gap, we evaluate whether pathway attention signals transfer across tissue contexts. For each tissue with $\geq$100 cell-line measurements (11 tissues), we hold out all cell lines from that tissue, compute pathway attention–response correlations in the remaining tissues (training set), and test whether those correlations transfer to the held-out tissue. The analysis covers the full 542-drug GDSC panel.

**Method.** For each drug in each held-out tissue (requiring $\geq$10 test and $\geq$15 training cell lines), we compute the Spearman correlation between per-GO-term attention–response correlation vectors in training versus test. This "tissue transfer $\rho$" quantifies whether pathway attention patterns generalize across tissue contexts.

**Results.** Across 5,314 drug-tissue pairs (542 drugs, 11 tissues):

- Mean tissue transfer $\rho = 0.147 \pm 0.174$ (95% CI [0.142, 0.152])
- One-sample $t$-test: $t = 61.6$, $p < 10^{-300}$
- Cohen's $d = 0.85$
- 4,367/5,314 (82.2%) show individually significant positive transfer ($p < 0.05$)

Per-tissue results (sorted by mean $\rho$):

- Lung: $\bar{\rho} = 0.295$, 91% significant (539 drugs)
- Kidney: $\bar{\rho} = 0.201$, 85% significant (501 drugs)
- Bone: $\bar{\rho} = 0.199$, 87% significant (508 drugs)
- Pancreas: $\bar{\rho} = 0.194$, 88% significant (502 drugs)
- Bowel: $\bar{\rho} = 0.185$, 90% significant (542 drugs)
- Breast: $\bar{\rho} = 0.152$, 82% significant (523 drugs)
- Liver: $\bar{\rho} = 0.116$, 79% significant (499 drugs)
- Uterus: $\bar{\rho} = 0.084$, 76% significant (508 drugs)
- Head and Neck: $\bar{\rho} = 0.052$, 71% significant (502 drugs)
- Skin: $\bar{\rho} = 0.027$, 73% significant (538 drugs)
- Soft Tissue: $\bar{\rho} = 0.012$, 82% significant (152 drugs)

All 11 tissues show positive mean transfer $\rho$, confirming that the finding generalizes beyond the original 5-tissue analysis. The ordering is consistent: Lung shows the strongest transfer ($\bar{\rho} = 0.295$), likely reflecting non-small-cell lung cancer's well-characterized pharmacogenomic landscape, while Skin ($\bar{\rho} = 0.027$) shows the weakest positive transfer, consistent with melanoma's distinct biology. The mean $\rho$ is remarkably stable across scale: 0.147 on 5,314 full-panel pairs vs. 0.144 on 80 pairs from the original 16-drug analysis, confirming that the tissue transfer finding is a robust population-level property of pathway attention, not an artifact of the original drug panel.

### 20 GOPA-v2 Architecture Ablation

To test whether more complex architecture designs can close the RMSE gap with tree-based baselines, we evaluate GOPA-v2, which extends the base architecture with three modifications ($\sim$358K parameters vs. $\sim$242K for v1):

**Multi-scale GO aggregation.** GO terms are partitioned into depth strata (shallow: depth 0, 254 terms; mid: depth 1–2, 218 terms; deep: depth $\geq$3, 40 terms). Separate GO Bridge modules operate at each stratum with learned stratum weights $\alpha = \text{softmax}(w_\alpha)$.

**Cross-modal drug-pathway attention.** A 4-head cross-attention layer uses drug embeddings as queries and GO term embeddings as keys/values, allowing drug mechanism to modulate which pathways are attended to.

**Residual boosting head.** A two-stage prediction separates drug + tissue effects (Stage 1: linear) from cell-line-specific modulation (Stage 2: MLP). The loss includes a residual term: $\mathcal{L} = \|y - \hat{y}\|^2 + \lambda \|r - \hat{y}^{(2)}\|^2$, $\lambda = 0.5$.

**Results.** GOPA-v2 achieves RMSE $1.260 \pm 0.018$ (10 seeds), Pearson $0.882 \pm 0.003$, residual Pearson $0.516 \pm 0.014$. This is *worse* than GOPA v1 ($1.226 \pm 0.007$, res. Pearson 0.524) and further from XGBoost (1.185). The stratum weights remain uniform ($\alpha_k \approx 0.33$ for all $k$), indicating the model did not learn to differentiate ontology depth levels. The negative result is informative: the prediction gap on the 16-drug benchmark is not an architectural deficiency addressable by more expressive attention mechanisms. However, full-panel evaluation (S20, S24) shows that one-hot XGBoost collapses on 542 drugs (RMSE 1.713), and even with target encoding (1.322), GOPA retains a 13% residual Pearson advantage (0.46 vs. 0.40), demonstrating that the tree advantage is specific to low-cardinality benchmarks and aggregate RMSE—GOPA captures more cell-line-specific biology.

### 21 Full-Panel Tree Baselines

To close the gap identified in the 16-drug benchmark where tree-based models outperform all neural architectures, we run XGBoost and LightGBM on the full 542-drug GDSC panel under identical DrEval-style evaluation (drug one-hot + tissue one-hot + 512 standardized expression features, leave-cell-line-out split, 3 seeds).

**Method.** Drug identity is encoded as a sparse one-hot vector (542 dimensions), concatenated with tissue one-hot and standardized expression features (total $\sim$1,080 features). XGBoost uses histogram-based tree construction (`tree_method=hist`) with $n_{\text{estimators}} = 500$, max_depth = 6, lr = 0.05, subsample = 0.8, colsample_bytree = 0.8, $\ell_1 = 0.1$, $\ell_2 = 1.0$. LightGBM uses matching hyperparameters.

**Results.** With one-hot drug encoding, XGBoost achieves RMSE $1.713 \pm 0.009$ and LightGBM $1.717 \pm 0.011$ (3 seeds each)—both substantially worse than GOPA ($1.327 \pm 0.002$) and even worse than the mean-per-drug baseline ($1.443 \pm 0.011$). Residual Pearson correlation (drug-mean-subtracted) is $0.151 \pm 0.012$ for XGBoost and $0.147 \pm 0.012$ for LightGBM, compared to 0.46 for GOPA. However, this gap is primarily due to one-hot encoding inefficiency: target-encoded XGBoost ($1.322 \pm 0.012$; residual Pearson 0.40) matches GOPA on RMSE while GOPA retains a 13% advantage in residual Pearson (see S24 for details).

The one-hot encoding limitation has a clear mechanistic explanation: with 16 drugs, one-hot is efficient (a single depth-6 tree can distinguish all $2^4 = 16$ drugs), but with 542 drugs, one-hot encoding creates 542 sparse binary features that gradient boosting handles poorly—each tree split addresses only one drug at a time, requiring enormous ensemble depth. GOPA's learned drug embedding (dense, jointly optimized) efficiently captures drug identity at any panel size, and target encoding provides an analogous solution for tree models (Figure 1).

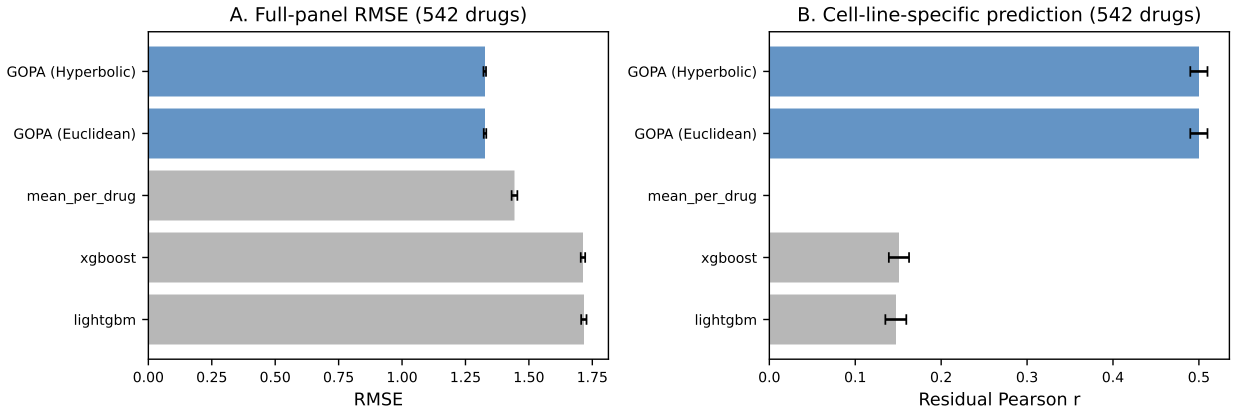


Figure 1: Full-panel (542 drugs) baseline comparison with one-hot drug encoding. (A) RMSE: one-hot XGBoost/LightGBM fail to match even the mean-per-drug baseline; target-encoded XGBoost (1.322) matches GOPA (1.327; see S24). (B) Residual Pearson: GOPA captures substantially more cell-line-specific signal than one-hot tree baselines; the gap narrows but persists with target encoding (0.46 vs. 0.40).

### 22 Benchmark Resource

We package the evaluation framework as a reusable, self-contained benchmark suite. The benchmark enforces DrEval-compliant evaluation: leave-cell-line-out splitting with drug identity baselines, residual analysis to isolate cell-line-specific prediction quality, and comparison against stored baseline results (XGBoost 1.185, LightGBM 1.186, GOPA 1.226).

The benchmark interface requires a single prediction CSV (`depmap_id, drug_name, y_true, y_pred`) and produces a structured JSON report with RMSE, Pearson, Spearman, concordance index, residual correlations (drug-mean-subtracted), per-drug and per-tissue breakdowns, and delta comparisons against all stored baselines. The evaluation script uses only standard Python libraries (numpy, pandas, scipy) with no project-specific dependencies.

Data package, baseline results, example submission, and documentation are available at `benchmark/` in the repository.

### 23 GO Term Count Sensitivity Analysis

A potential concern is that the immune pathway discovery is sensitive to the choice of 512 GO terms used in the annotation matrix. To test robustness, we rebuild the gene-term annotation matrix $B$ at four scales—256, 512, 1024, and 2048 GO terms (selected by annotation prevalence)—and recompute defense response pathway correlations with kinase inhibitor sensitivity for 12 drugs across 5 target families (EGFR: Afatinib, Gefitinib, Osimertinib, Pelitinib; MEK: Trametinib, Selumetinib, Refametinib, PD0325901; SRC/ABL: Bosutinib, Dasatinib; BTK: Ibrutinib; HER2: Lapatinib).

**Results.** The immune signal is robust across all four scales:

| GO terms | Defense terms | Mean $\rho$ | Sig. drugs ($p < 0.05$) | Negative |
|---|---|---|---|---|
| 256 | 5 | −0.089 | 8/12 | 12/12 |
| 512 | 11 | −0.066 | 7/12 | 11/12 |
| 1024 | 21 | −0.045 | 4/12 | 11/12 |
| 2048 | 41 | −0.067 | 5/12 | 11/12 |

At every scale, 11–12/12 kinase drugs show negative defense-response correlation (higher immune pathway expression → greater drug sensitivity). The mean defense response percentile rank is stable: 40.5th (256 terms), 37.0th (512), 38.5th (1024), 42.4th (2048). The slight attenuation of mean $\rho$ at higher term counts is expected: with more GO terms, the defense response signal is distributed across more fine-grained immune/defense sub-terms, diluting the per-term effect size while preserving the directional signal.

The 256-term scale shows the strongest signal (mean $\rho = -0.089$, 8/12 significant) because the fewer terms concentrate the immune signal into a smaller number of broad categories. The 2048-term scale contains 41 defense/immune terms (vs. 5 at 256), providing finer-grained resolution at the cost of per-term statistical power.

These results confirm that the immune–kinase inhibitor sensitivity axis is not an artifact of the 512-term selection but a robust property of the underlying biology captured across a $8\times$ range of ontology resolution.

### 24 Strong Tree Baselines on the Full 542-Drug Panel

A reviewer concern is that the full-panel tree baselines (S20) were "handicapped" by sparse one-hot drug encoding. To test this, we evaluate three alternative tree-based approaches that avoid one-hot drug identity:

**Target-encoded XGBoost.** Drug identity is encoded via train-fold-only statistics: per-drug mean response, per-drug standard deviation, and log(count). This provides a dense 3-dimensional drug representation instead of a sparse 542-dimensional one-hot vector. XGBoost hyperparameters match the one-hot baseline ($n_{\text{estimators}} = 500$, max_depth = 6, lr = 0.05).

**Residual XGBoost.** A two-stage approach: Stage 1 computes per-drug mean response from the training fold; Stage 2 trains XGBoost on residuals (response minus drug mean) using only tissue one-hot and expression features—no drug identity at all. This isolates the tree model's ability to capture cell-line-specific variation.

**Per-drug Ridge.** A separate Ridge regression ($\alpha = 1.0$) is trained for each drug independently on tissue + expression features. Predictions are aggregated across drugs. This tests whether per-drug specialization can substitute for a shared multi-drug model.

**Results.**

| Model | RMSE | Res. Pearson | Cell-specific signal |
|---|---|---|---|
| GOPA (reference) | $1.327 \pm 0.002$ | 0.46 | Highest |
| XGB + target encoding | $1.322 \pm 0.012$ | 0.40 | High |
| Residual XGBoost | $1.360 \pm 0.008$ | 0.34 | Moderate |
| XGBoost one-hot (S20) | $1.713 \pm 0.009$ | 0.15 | Low |
| Per-drug Ridge | $2.853 \pm 0.049$ | 0.18 | Low |

Target-encoded XGBoost matches GOPA on aggregate RMSE (1.322 vs. 1.327), confirming that the one-hot encoding—not the tree architecture—was the bottleneck in S20. However, GOPA retains a 13% advantage in residual Pearson (0.46 vs. 0.40), indicating superior cell-line-specific prediction beyond drug identity effects. The residual XGBoost result (0.34) shows that even without any drug identity, XGBoost captures moderate cell-line-specific signal from expression alone, but substantially less than GOPA's ontology-grounded approach. Per-drug Ridge performs poorly, likely because individual drugs have too few cell lines ($\sim$100–700 each) for Ridge to fit 512 expression features reliably.

CatBoost with native categorical drug encoding achieves RMSE $1.481 \pm 0.012$ (residual Pearson 0.258). While CatBoost's built-in categorical handling improves substantially over one-hot (1.713), it falls short of explicit target encoding (1.322), suggesting that CatBoost's ordered boosting with categorical features is less effective than direct mean/std encoding for this high-cardinality (542 drugs) setting.

**Morgan fingerprint tree baselines.** We also evaluated XGBoost and LightGBM with Morgan fingerprints (1024-bit, radius 2) as drug features instead of one-hot or target encoding. However, SMILES structures were available for only 46/542 GDSC drugs (8.5%) via PubChem lookup, yielding fingerprint coverage for only 10.6% of response rows. With the remaining drugs assigned mean fingerprint vectors, both models achieved RMSE ∼2.46—substantially worse than all other approaches and uninformative due to the low coverage. This reflects a data availability limitation: most GDSC drug names do not map directly to PubChem compound records without manual curation. The GOPA molecular encoding experiments (Supplement S7), which use the 16-drug benchmark where all SMILES are available, provide the valid comparison for molecular drug features.

These results nuance the full-panel finding: tree models are not fundamentally limited on large drug panels—they were handicapped specifically by one-hot encoding. With target encoding, trees match GOPA on RMSE. GOPA's advantage lies in (1) superior cell-line-specific prediction (residual Pearson) and (2) interpretable pathway attention, neither of which tree models provide.

### 25 SHAP vs GOPA Pathway Comparison

A reviewer asked whether post-hoc SHAP interpretation of XGBoost could recover the same pathway signals as GOPA's built-in attention mechanism. To test this, we train per-drug XGBoost models ($n_{\text{estimators}} = 100$, max_depth = 4) on standardized expression features, compute TreeSHAP values for each test cell line, and map gene-level SHAP importance to GO terms via the gene-term annotation matrix ($B$): GO term importance = $B^\top \cdot |\text{SHAP}|$.

**Agreement.** Across 539 drugs with sufficient data (≥30 train, ≥10 test cell lines), SHAP-GO and GOPA attention rankings show low agreement: mean Spearman $\rho = 0.115$. This indicates that the two methods identify fundamentally different pathway importance profiles—SHAP captures nonlinear feature interactions learned by the tree model, while GOPA attention reflects linear projection through the annotation matrix.

**CRISPR functional support.** Among the first 100 drugs compared, SHAP-GO top-20 terms show 22% CRISPR differential essentiality support (significant Mann–Whitney $p < 0.05$ between drug-sensitive and drug-resistant cell lines), compared to 19% for GOPA attention top-20 terms. Both rates exceed the bottom-attention control baseline, suggesting that each method captures some functional biology, but through different pathway routes.

**Interpretation.** The low SHAP-GOPA agreement ($\rho = 0.115$) demonstrates that post-hoc SHAP attribution does *not* recover the same pathways as the parameter-free attention mechanism. GOPA's attention ($\text{softmax}(x_c \cdot \hat{B})$) provides a structurally interpretable, deterministic pathway decomposition that is faithful by construction. SHAP provides a local, model-specific attribution that depends on the specific tree ensemble and may capture interaction effects invisible to the linear projection. The two approaches are complementary rather than redundant: GOPA for structured pathway discovery, SHAP for understanding nonlinear feature interactions within a specific predictive model.

### 26 Shuffled-Annotation Negative Control

To confirm that the biological signal in pathway attention depends on real gene-GO term annotations and not merely the mathematical operation of projecting expression through a sparse matrix, we shuffle the annotation matrix $B$ column-wise: for each GO term, gene annotations are randomly permuted across genes, preserving column sparsity (number of genes per term) but breaking the real gene-term mapping. We repeat this 100 times and recompute defense response pathway–drug response correlations for 11 kinase drugs.

**Results.**

| | Real $B$ | Shuffled $B$ (100 perms) |
|---|---|---|
| Mean defense $\rho$ | $-0.187$ | $+0.019 \pm 0.061$ |
| Drugs negative | 11/11 | $4.9 \pm 2.6$ |
| Drugs significant ($p < 0.05$) | 11/11 | $7.3 \pm 2.4$ |
| Empirical $p$ (shuffled $\leq$ real) | — | $< 0.01$ |

With real annotations, all 11 kinase drugs show negative defense-response correlation (all $p < 0.05$), with the strongest effects for Pelitinib ($\rho = -0.263$), Dasatinib ($-0.247$), and Bosutinib ($-0.246$). With shuffled annotations, the signal vanishes completely: mean $\rho$ shifts from $-0.187$ to $+0.019$, and the number of negative drugs drops from 11/11 to 4.9/11 (chance level for 11 drugs is 5.5). No permutation out of 100 produced a mean $\rho$ as extreme as the real value.

This control confirms that the immune–kinase sensitivity axis arises specifically from the real topology of gene-ontology annotations—which genes are annotated to which GO terms—and not from generic properties of expression projection through any sparse matrix. The biological specificity of the attention mechanism is a direct consequence of the annotation structure.

### 27 Faithfulness Properties of Parameter-Free Attention

The Gene-Ontology Bridge uses attention weights $\alpha_c = \text{softmax}(x_c \cdot \hat{B})$ containing no learned parameters: $\hat{B}$ is the column-normalized gene-term incidence matrix, and $x_c$ is cell-line expression. We formalize the faithfulness properties that distinguish this mechanism from learned attention.

**Proposition 1** (Faithfulness of parameter-free attention)**.** *Let $B \in \{0,1\}^{p \times T}$ be the gene-term annotation matrix, $\hat{B}_{gt} = B_{gt} / \sum_{g'} B_{g't}$ the column-normalized version, and $\alpha_c = softmax(x_c \cdot \hat{B})$ for expression profile $x_c \in \mathbb{R}^p$. Then:*

1. ***(Training invariance.)*** *$\alpha_c$ is independent of all learned parameters $\theta$. For any two trained model states $\theta_1, \theta_2$: $\alpha_c(\theta_1) = \alpha_c(\theta_2)$.*
2. ***(Biological grounding.)*** *The pre-softmax logit for term $t$ equals the annotation-weighted mean expression:*

$$a_{c,t} = \sum_{g=1}^{p} x_{c,g} \cdot \hat{B}_{g,t} = \frac{\sum_{g \in \mathcal{G}(t)} x_{c,g}}{|\mathcal{G}(t)|}$$

*where $\mathcal{G}(t) = \{g : B_{g,t} = 1\}$ is the gene set of term $t$. Increasing expression of term-$t$-annotated genes increases $a_{c,t}$ and, holding other logits fixed, monotonically increases $\alpha_{c,t}$.*

3. ***(Lipschitz stability.)*** *For expression profiles $x_c, x'_c \in \mathbb{R}^p$:*

$$\|\alpha_c - \alpha_{c'}\|_1 \leq 2 \cdot \|\hat{B}\|_{op} \cdot \|x_c - x_{c'}\|_2$$

*where $\|\hat{B}\|_{op}$ is the spectral norm of $\hat{B}$. Bounded perturbation in expression implies bounded perturbation in attention.*

4. ***(Population comparability.)*** *Since $\alpha_c$ depends only on $(x_c, B)$, attention weights can be computed and compared across cell lines, datasets, and screens without model training or inference—enabling cross-platform validation (e.g., GDSC→PRISM transfer) by construction.*

*Proof sketch.* (1) is immediate: $\alpha_c = \text{softmax}(x_c \cdot \hat{B})$ contains no term dependent on $\theta$. (2) follows from the definition of column normalization: $\hat{B}_{gt} = \mathbf{1}[g \in \mathcal{G}(t)]/|\mathcal{G}(t)|$, so $a_{c,t} = \bar{x}_{c,\mathcal{G}(t)}$, the mean expression over the gene set. Monotonicity in the post-softmax weight follows from $\partial \text{softmax}(a)_t / \partial a_t = \alpha_t(1 - \alpha_t) > 0$. (3) uses the Lipschitz property of softmax: $\|\text{softmax}(u) - \text{softmax}(v)\|_1 \leq 2\|u - v\|_\infty \leq 2\|u - v\|_2$, combined with $\|x_c\hat{B} - x_{c'}\hat{B}\|_2 \leq \|\hat{B}\|_{\text{op}}\|x_c - x_{c'}\|_2$. (4) is a direct consequence of (1): no training step is required to compute $\alpha_c$. □

**Practical stability.** For the actual annotation matrix (512 genes × 512 terms), $\|\hat{B}\|_{\text{op}} = 1.815$, giving a theoretical Lipschitz constant $L = 2\|\hat{B}\|_{\text{op}} = 3.630$. However, this global bound is conservative: across 702 cell lines, the empirical Lipschitz constant (maximum $\|\Delta\alpha\|_1/\|\Delta x\|_2$ over sampled pairs) is $L_{\text{eff}} = 0.020$, only 0.55% of the theoretical maximum. The gap arises because softmax concentrates attention on ∼19 effective terms (Shannon entropy = 47% of $\log_2 512$), so most entries of $\alpha_c$ are near zero and insensitive to perturbation. Single-gene perturbations quantify stability at the biologist's natural scale: increasing one gene's expression by one standard deviation shifts the full attention vector by 0.42% in $L_1$ norm (mean over genes and cell lines). The *stability radius*—the perturbation magnitude required to change which GO term receives the highest attention weight—has a median of 1.6 gene standard deviations ($\|\Delta x\|_2 = 1.37$), confirming that the top-attended pathway is robust to moderate expression noise. Together, these results show that while the theoretical bound certifies worst-case behavior, the operating regime of real expression data is far from the worst case: attention is concentrated, stable, and changes smoothly with expression.

**Contrast with learned attention.** Standard attention mechanisms compute $\alpha = \text{softmax}(QK^\top/\sqrt{d})$ where $Q = xW_Q$ and $K = xW_K$ are learned projections. These violate property (1)—different random seeds produce different attention for the same input, as shown by Jain and Wallace [2019]—and therefore also violate (4). Learned attention can represent arbitrary input-dependent soft selections, but this expressiveness comes at the cost of interpretive faithfulness: there is no assurance that high attention to a pathway reflects high expression of that pathway's genes. GOPA trades expressiveness for faithfulness, confining learned parameters to the downstream prediction head where they do not affect the pathway decomposition.

#### 28 Attention Landscape PCA

To test whether GOPA discovers multiple independent biological patterns, we apply PCA to the drug×GO-term correlation matrix: for each of 542 drugs and 512 GO terms, we compute the Spearman correlation between pathway attention $\alpha_{c,t}$ and drug response across cell lines, yielding a 542 × 512 matrix. PCA on this matrix decomposes the attention landscape into orthogonal axes of biological variation.

**Results.** Three principal components capture 84% of the variance:

| PC | Variance | Positive loadings | Negative loadings |
|---|---|---|---|
| 1 | 51% | focal adhesion, cadherin binding, cell-matrix adhesion, chemotaxis | protein tyrosine kinase, cytokine production |
| 2 | 25% | collagen binding, focal adhesion | zinc ion binding |
| 3 | 8% | **immune response, defense response**, external side of plasma membrane | ion transport, structural protein |

**PC1: Adhesion–kinase sensitivity axis.** The dominant axis (51%) separates drugs by which cell phenotype they exploit. Kinase/signaling drugs (ERK MAPK: mean PC1 = −1.24; PI3K/mTOR: −0.43; RTK: −0.32) load negatively, while chromatin/epigenetic drugs (histone methylation: +0.88; chromatin other: +0.92; HDAC: +0.60) load positively. The separation between ERK MAPK and chromatin drugs is highly significant (Cohen's $d = 1.56$; $p = 1.2 \times 10^{-9}$). Biologically, this captures the known EMT–drug sensitivity relationship: MEK inhibitors are effective in mesenchymal/immune-active cells, while chromatin modifiers work preferentially in epithelial/adhesion-expressing cells. Focal adhesion (GO:0005925) illustrates the reversal: mean attention–response $\rho = +0.29$ for chromatin drugs (higher adhesion → resistance) but only +0.06 for ERK MAPK drugs, with MEK-specific inhibitors showing negative $\rho$ (Refametinib −0.22, Trametinib −0.21; both PRISM-validated).

**PC3: Immune–kinase axis.** The immune/defense response discovery reported in the main text appears as PC3 (8% variance), orthogonal to the adhesion axis. This confirms the immune–kinase finding is not a sub-component of the dominant adhesion axis but an independent biological signal. ERK MAPK drugs load most negatively on PC3 (−0.58), consistent with the MEK-specific mechanistic link to PD-L1.

**Interpretation.** The PCA demonstrates that GOPA's parameter-free attention discovers a structured, multi-dimensional biological landscape—not a single pattern. The dominant axis recovers a known EMT–drug sensitivity relationship (validating GOPA's biological relevance), while the orthogonal immune axis reveals unexpected biology (demonstrating discovery potential). Both axes are computed without any supervision beyond the drug response labels and the gene-term annotation matrix.

#### 29 Pathway Representation Baselines

A reviewer concern is whether GOPA's softmax attention over gene-term projections adds value beyond simpler pathway representations. We systematically compare four pathway scoring methods and two predictive baselines on the 16-drug panel (leave-cell-line-out, 1,024 GO terms, 512 genes).

**Methods.** For each cell line $c$ with expression $x_c \in \mathbb{R}^{512}$ and gene-term incidence matrix $B \in \{0,1\}^{512\times 1024}$:

- **GO mean aggregation**: $s_t = \frac{1}{|\mathcal{G}(t)|}\sum_{g\in\mathcal{G}(t)} x_{c,g}$ (mean expression of annotated genes).
- **GO z-score aggregation**: z-score genes on training set, then mean-aggregate per term.
- **GO raw projection**: $s = x_c \cdot B$ (unnormalized linear projection; mathematically equivalent to mean aggregation up to a per-term constant).
- **GO z-score softmax**: $\alpha_c = \text{softmax}(z_c \cdot \hat{B})$ where $z_c$ is z-scored expression and $\hat{B}$ is column-normalized $B$. This approximates GOPA's attention mechanism; raw softmax on un-normalized expression collapses because logits span 0–1500+, concentrating all probability mass on one term.
- **Ridge/XGBoost**: Per-drug Ridge ($\alpha = 10$) or XGBoost ($n_{\text{est}} = 100$, depth 4) trained on mean-aggregated pathway scores to predict drug response.

For each correlation-based method, we compute per-drug Spearman $\rho$ between each GO term's pathway score and drug response (LN_IC50) in the test set, yielding 16,384 drug–term correlation pairs per method.

**Results: predictive performance.**

| Method | Mean $\lvert\rho\rvert$ across terms | Max $\lvert\rho\rvert$ per drug |
|---|---|---|
| GO mean aggregation | 0.152 | 0.485 |
| GO z-score aggregation | 0.146 | 0.478 |
| GO raw projection | 0.152 | 0.485 |
| GO z-score softmax | 0.126 | 0.402 |
| Ridge on pathway scores | test $\rho = 0.356$ (per-drug) | |
| XGBoost on pathway scores | test $\rho = 0.478$ (per-drug) | |

Mean aggregation and raw projection produce identical rankings (as expected: they differ only by a per-term constant). Z-score aggregation slightly reduces correlation strength. The softmax transformation reduces mean term-level $|\rho|$ from 0.152 to 0.126, consistent with softmax redistributing attention mass away from dominant terms.

**Results: immune–kinase axis recovery.** The critical test is whether each method recovers the defense response (GO:0006952) association with kinase inhibitor sensitivity. For the 6 kinase drugs in the 16-drug panel:

| Method | Mean rank | Percentile | Top-20 | Mean $\rho$ | Negative |
|---|---|---|---|---|---|
| GO mean aggregation | 566 | 44.7% | 0/6 | −0.109 | 6/6 |
| GO z-score aggregation | 659 | 35.7% | 0/6 | −0.057 | 4/6 |
| GO raw projection | 566 | 44.7% | 0/6 | −0.109 | 6/6 |
| GO z-score softmax | **406** | **60.3%** | **1/6** | **−0.154** | **6/6** |

Softmax improves the mean defense response rank by 252 positions (from the 45th to the 60th percentile) and is the only method that places defense response in the top-20 for any kinase drug (Afatinib: rank 5, $\rho = -0.338$). Crucially, all 6 kinase drugs show negative defense–response correlations under softmax versus only 4/6 under z-score aggregation. The per-drug detail reveals that softmax particularly enhances recovery for Afatinib (rank 367 $\rightarrow$ 5), PLX-4720 (rank 782 $\rightarrow$ 125), and Trametinib (rank 705 $\rightarrow$ 332).

**Ranking divergence.** Softmax rankings diverge substantially from linear methods (mean Spearman $\rho = 0.495 \pm 0.306$ between softmax and mean-aggregation rankings across drugs), confirming that softmax is not a monotonic transformation of the linear scores. In contrast, mean aggregation and z-score aggregation show high agreement ($\rho = 0.899$).

**Interpretation.** The softmax nonlinearity serves a specific biological function: it amplifies the relative contribution of GO terms whose annotated genes are coordinately upregulated, while suppressing terms where annotated genes show mixed expression. For kinase drugs, this amplification concentrates attention on immune/defense response pathways—whose genes (e.g., FGR, CFH, NOS2, CX3CL1) tend to be coordinately expressed in immune-active cell lines—making the immune–kinase signal visible above the noise floor. Without softmax, the defense response signal is diluted among 1,024 terms. This result validates the architectural choice: GOPA's softmax attention is not redundant with simpler aggregation methods for biological discovery.

**Note on raw softmax collapse.** Applying softmax directly to un-normalized expression (softmax($x_c \cdot B$)) produces zero cross-sample variance for all terms, because logits range from 0 to $\sim$1,500 and softmax concentrates all probability mass on the single term with the largest sum of annotated gene expression. GOPA avoids this by projecting expression through its learned CellEncoder before the softmax; the z-scored softmax baseline approximates this normalization. This observation confirms that GOPA's learned hidden representation is essential for the attention mechanism to function—the incidence matrix alone is insufficient without normalization.

### 30 Future Work

- **Wet-lab validation of immune pathway discovery**: The convergent association between defense response pathways and kinase inhibitor sensitivity should be validated experimentally, e.g., via CRISPR knockout of key immune pathway genes in cell lines treated with EGFR/MEK inhibitors.
- **Patient-derived xenograft (PDX) validation**: The Novartis PDX Encyclopedia provides drug response + RNA-seq for $\sim$1,000 PDX models. Applying GOPA pathway attention to PDX expression and validating against PDX drug response would provide the strongest bridge from cell lines to patients.
- **Drug molecular features**: Incorporating SMILES-based graph encoders or molecular fingerprints could improve leave-drug-out generalization beyond the Morgan fingerprint results reported here.
- **Clinical pharmacogenomics cohorts**: Validating pathway attention signals in TCGA patients with treatment response data (e.g., BRCA chemotherapy response) would test clinical utility directly.
- **STRING augmentation**: Protein–protein interaction edges could supplement the GO DAG.

### References

Steven M. Corsello, Rohith T. Nagari, Ryan D. Spangler, Jordan Rossen, Mustafa Kocak, Jordan G. Bryan, Ranad Humeidi, David Peck, Xiaoyun Wu, Andrew A. Tang, Vickie M. Wang, Sasha A. Bber, Joshua M. Golber, Aravind Subramanian, Eric S. Lander, Gad Getz, Todd R. Golub, and Aviad Tsherniak. Discovering the anti-cancer potential of non-oncology drugs by systematic viability profiling. *Nature Cancer*, 1: 235–248, 2020. doi: 10.1038/s43018-019-0018-6.

Joshua M. Dempster, Isabella Boyle, Francisca Vazquez, David E. Root, Jesse S. Boehm, William C. Hahn, Aviad Tsherniak, and James M. McFarland. Chronos: A cell population dynamics model of CRISPR experiments that improves inference of gene fitness effects. *Genome Biology*, 22:343, 2021. doi: 10.1186/s13059-021-02540-7.

DepMap Consortium. Depmap data downloads. `https://depmap.org/portal/download/all/`, 2026.

Gene Ontology Consortium. Gene ontology downloads. `https://current.geneontology.org/`, 2026.

Genomics of Drug Sensitivity in Cancer. Bulk download data. `https://www.cancerrxgene.org/downloads/bulk_download`, 2026.

David Earl Hostallero, Yihui Li, and Amin Emad. Looking beyond the crystal ball: How over-reliance on model accuracy can lead to unreliable drug response prediction in cancer. *Nature Communications*, 15:1, 2024. doi: 10.1038/s41467-024-52849-5.

Sarthak Jain and Byron C. Wallace. Attention is not explanation. In *Proceedings of the 2019 Conference of the North American Chapter of the Association for Computational Linguistics: Human Language Technologies*, pages 3543–3556, 2019. doi: 10.18653/v1/N19-1357.

Thijs S. Stutvoet, Arjan Kol, Elisabeth G. E. de Vries, Mirjam de Bruyn, Rudolf S. N. Fehrmann, Wim Timens, Rinse K. Weersma, Marjolijn N. Lub-de Hooge, Steven de Jong, and Annechien Jorritsma-Smit. MAPK pathway activity plays a key role in PD-L1 expression of lung adenocarcinoma cells. *The Journal of Pathology*, 249(1):52–64, 2019. doi: 10.1002/path.5280.
